# KozakExplorer: an interactive framework for genome-wide Kozak sequence analysis

**DOI:** 10.64898/2026.06.25.734688

**Authors:** Thomas Cokelaer, Ana Maria Santi, Juliana Pipoli da Fonseca, Gerald F. Späth

## Abstract

Translation initiation signals shape gene expression across all domains of life. In eukaryotes, nucleotide constraints surrounding the start codon are commonly described by the Kozak Consensus Sequence (KCS), whereas in bacteria and archaea, initiation frequently involves Shine–Dalgarno ribosome-binding motifs. Although these signals have been extensively characterized in model organisms, their large-scale diversity and evolutionary distribution remain incompletely explored.

We present KozakExplorer, a reproducible framework for quantitative and comparative analysis of translation initiation contexts from genome assemblies and annotations. The software performs strand-aware extraction of start codon environments from FASTA and GFF3 files and applies information-theoretic metrics—including Kullback–Leibler (KL) divergence and information content (IC)—to measure positional nucleotide constraints relative to a background model. Derived summary statistics (Kozak Strength Index [KSI], maximum information content, peak position) convert motif patterns into interpretable per-genome signatures suitable for cross-species comparison. Our primary analysis covers 2,282 eukaryotic reference genomes, producing a standardized dataset of translation initiation metrics. Dimensionality reduction via t-SNE on per-position KL divergence, information content, and motif nucleotide frequencies reveals a structured eukaryotic KCS landscape with kingdom-level clustering and continuous variation in signal strength. A dedicated case study of 216 Apicomplexa genomes shows genus-level structure consistent with host range and phylogeny. An extended analysis across 25,344 reference genomes (22,253 bacteria, 809 archaea) places eukaryotic patterns in a global comparative framework, revealing transitions between sharply localized Kozak motifs and distributed Shine–Dalgarno-type signatures.

Implemented within the open-source Sequana ecosystem, KozakExplorer is distributed as a Python module and an interactive web application that accepts local annotated assemblies, GenBank records, or NCBI RefSeq accessions, and exports all computed metrics, embeddings, and coordinates for downstream comparative and evolutionary genomics.

**Availability:** Implemented in Python within the Sequana framework. Source code available at https://github.com/sequana/sequana and https://github.com/sequana/webapp_kozak.

## 1 Introduction

Translation initiation is a central regulatory step of gene expression in all domains of life [1]. In eukaryotes, recognition of the start codon by the scanning ribosome is strongly influenced by the nucleotide context flanking the AUG codon, a dependence first characterized by Kozak [2, 3]. Three related terms are used to describe this context, and we distinguish them throughout this work. The *Kozak sequence* is the gene-specific nucleotide context surrounding the start codon of an individual coding sequence. The *Kozak consensus sequence* (KCS) is the species-level summary of these sequences, in which each position is reduced to a single consensus letter (for example, a purine at position −3) [3, 4]. The rule for assigning the consensus letter at each position varies between studies, from the simplest majority base to the stricter Cavener convention [5]. A complementary, coarser scheme instead grades each context by its two functionally dominant positions—a purine at −3 and a *G* at +4—as *strong*, *adequate* (moderate), or *weak* [3, 6, 7]. The *Kozak motif* retains the full per-position nucleotide-frequency distribution—as a position frequency matrix or sequence logo—rather than collapsing it to a single consensus letter, and therefore preserves the strength of the constraint at each position. Canonical KCS strings such as GCC(A/G)CCAUGG were initially characterized in vertebrates, yet subsequent studies have demonstrated substantial interspecies variability, particularly among unicellular eukaryotes and parasitic protists [8, 7]. The Kozak motif, by contrast, preserves both the strength and the gene-to-gene variation that consensus reduction discards.

We further distinguish the *translation initiation site* (TIS), the specific codon at which translation begins, from the surrounding nucleotide context that modulates its recognition by the ribosome. We refer to this context—the window of nucleotides flanking the start codon—as the *initiation region* (IR) [9], a domain-neutral term that subsumes both the eukaryotic Kozak sequence and the bacterial and archaeal Shine–Dalgarno region [10]. In this work, rather than predicting the site, we start from TIS coordinates already annotated in reference genomes and quantify the positional nucleotide signal within the initiation region.

The functional impact of this context is striking: recent sort-seq libraries spanning the six nucleotides preceding the AUG show that Kozak variation alone can tune steady-state protein abundance over a ∼100-fold range and can identify germline variants with expression-reducing substitutions in cancer-associated genes [11]. Beyond steady-state output, Kozak strength also governs leaky scanning, allowing alternative downstream or upstream ORFs (uORFs) to be selected when the primary start is weak, thereby linking initiation context to regulatory and disease-relevant translational programs [12, 1]. Understanding this variability is therefore essential for elucidating translational regulation, improving genome annotation, and interpreting evolutionary adaptations in gene expression.

In bacteria and archaea, translation initiation relies on a distinct but functionally analogous signal: the Shine–Dalgarno sequence, a purine-rich motif typically located a few nucleotides upstream of the start codon [10]. Rather than encoding context around the AUG itself, the Shine–Dalgarno signal works by base-pairing with the ribosome to align the start codon for initiation [9, 13]. Despite mechanistic differences between scanning-dependent eukaryotic initiation [4] and bacterial or archaeal ribosome-binding-site recognition, both systems encode positional nucleotide constraints within the initiation region that can be quantitatively measured. From this perspective, Kozak and Shine–Dalgarno sequences represent alternative evolutionary solutions to the same functional requirement: accurate positioning of the start codon during translation initiation [14].

While consensus motifs provide a qualitative description of the initiation region, they do not offer a scalable framework for quantitative comparison across thousands of genomes: each genome is reduced to a single fixed-letter string that discards both the per-position constraint strength and the gene-to-gene variation, yielding no scalar summary that can be ranked, correlated, or embedded. Two genomes sharing the same consensus string may nonetheless differ markedly in the strength and localization of their initiation signal. With over 25,000 reference genomes now available in RefSeq spanning all three domains of life, there is an opportunity to investigate translation initiation architecture at unprecedented phylogenetic breadth. However, systematic, reproducible, and interactive tools capable of extracting, quantifying, and comparing initiation regions across diverse taxa remain limited. Existing resources address adjacent problems but not this one: sequence-logo tools such as WebLogo [15] render per-position frequencies for a single alignment without genome-scale aggregation; TIS-prediction tools such as NetStart 2.0 [16] focus on discriminating functional starts from spurious AUG codons rather than quantifying signal architecture; and prior surveys, though often substantial in their own scope—from hundreds of vertebrate mRNAs [3] and deep single-organism analyses of *Escherichia coli* ribosome-binding sites [9] to dozens of eukaryotic genomes [8] and broad reanalyses of published initiation-context data [7]—characterize the consensus qualitatively. None provides a scalar, per-genome summary of initiation signal architecture that is reproducible and comparable across the full breadth of RefSeq.

To address this need, we developed **KozakExplorer**, a modular framework integrated within the Sequana ecosystem [17] for large-scale analysis of initiation regions. Although named for the eukaryotic context that motivated its development, KozakExplorer is organism-agnostic: it applies the same positional analysis to any annotated genome, regardless of the underlying initiation mechanism. By taking annotated TIS coordinates as input, KozakExplorer complements rather than replaces TIS predictors, providing the kind of cross-species sequence statistics that such predictors rely on for training and benchmarking.

The framework combines strand-aware extraction of annotated start codon contexts from FASTA and GFF3 files with information-theoretic quantification of positional constraints. In addition to classical sequence logos, KozakExplorer computes Kullback–Leibler (KL) divergence relative to multiple background models and information content (IC) per position, each capturing complementary aspects of nucleotide constraints, along with derived scalar summaries (peak position, motif concentration) characterizing signal strength and localization. Our analysis charts a eukaryotic Kozak landscape across kingdoms, investigates the Apicomplexa clade in depth to uncover genus-level structure, and concludes with a cross-kingdom embedding that places Kozak and Shine–Dalgarno signals on a single continuum.

## 2 Materials and Methods

### 2.1 Software stack: sequana.kozak and webapp_kozak

The analyses presented in this work rely on a two-layer software stack distributed within the Sequana ecosystem [17]. First, the user-facing layer is webapp_kozak (https://github.com/sequana/webapp_kozak), an in-teractive Streamlit application (v1.58; https://streamlit.io) whose landing page lets a user compute translation initiation metrics from a single genome on demand. The inputs accepted on this page are a local FASTA + GFF3 pair, a GenBank accession, or an NCBI RefSeq accession. RefSeq assemblies are retrieved automatically through the NCBI Datasets command-line interface, while GenBank accessions are fetched through the bioservices Python pack-age [18], which provides a programmatic wrapper around the Entrez efetch endpoint. For the chosen genome, the application extracts Kozak contexts, computes per-position nucleotide frequencies and information-theoretic profiles, and renders a sequence logo, KL profile, IC profile, and chi-square statistics in real time. Window sizes, start-codon filters, and pandas DataFrame query-style row filters [19] are user-configurable. All results are exportable as CSV for downstream integration. The metrics computed by the interactive page are defined in the following subsections.

Second, underneath the web application sits the `kozak` module of the `Sequana` Python library (https://github.com/sequana/sequana), which provides all primitives used throughout the study: context extraction, position frequency matrix (PFM) and information-theoretic metric computation, and motif visualization. The same `sequana. kozak` primitives are also driven by a Snakemake [20] pipeline (described in the Automated analysis workflow sub-section) used to precompute the multi-genome corpus on which the comparative landscapes of this study are built.

### 2.2 Extraction of translation initiation contexts

For the selected genome, the application parses the GFF3 annotation to enumerate CDS features and locate every annotated start codon on the matching FASTA assembly. A nucleotide window—the initiation region—is then extracted around each start. The upstream span (default 9 nt, covering positions −9 to −1 and thus encompassing the canonical 6-base Kozak window) and the downstream span (default 6 nt) are user-configurable in the web application and in the underlying Python module, so that the user can adjust the analysis window to the organism or hypothesis at hand. Note that the interactive web app uses a narrower default window than the batch pipeline; the window bounds used for the reference corpus are fixed to [−20, +6] as described in the Automated analysis workflow subsection.

When a CDS is annotated across multiple exons, only the most 5’ (start-codon-containing) exon is used: extraction does not cross intron boundaries. For bacterial and archaeal genomes, which lack intron-spliced isoforms, each CDS feature contributes a single context and the isoform-handling step is a no-op.

For genes with multiple annotated transcript isoforms, each isoform is treated independently via grouping on the GFF3 Parent attribute (transcript identifier). Concretely, the `_collapse_to_first_cds` routine retains, for every transcript, the single CDS row whose coordinate is 5’-most on the annotated strand (smallest start on the + strand, largest stop on the − strand); subsequent CDS rows of the same transcript correspond to internal exon boundaries and are discarded, since they do not carry a biological start codon. Consequently, a gene with *n* annotated transcript variants contributes *n* start codons to the analysis rather than a single representative or longest-transcript pick. This design preserves isoform-specific Kozak diversity, as alternative transcripts may exhibit distinct 5’ UTRs, distinct start codons, and distinct translational efficiencies. The trade-off is that, in genomes with fragmented or over-annotated GFF3 files—typical of non-model organisms—high isoform counts can inflate the per-gene contribution to genome-wide statistics (KSI, entropy, information content) and, in turn, influence the t-SNE embeddings. Users analyzing newly assembled or legacy eukaryotic genomes should therefore inspect the per-gene transcript count distribution before interpreting derived metrics; the `collapse_first_cds` parameter of the `Kozak` class can also be disabled to recover the legacy one-row-per-CDS behavior for benchmarking purposes.

Strand orientation is fully respected: for genes annotated on the reverse strand, the relevant exon coordinates are taken from the GFF3 and the extracted sequence is reverse-complemented so that all contexts share a consistent orientation relative to the start codon.

By default, only canonical ATG start codons are retained. This is a conservative choice that may miss alternative initiation signals; the user can opt into extracting other start codons through the application’s filter controls or the corresponding parameter of the `Kozak` class.

As a sanity check, the interactive page also displays the empirical start-codon distribution of the annotation (counts of ATG, GTG, TTG, etc.). This plot is useful to confirm that ATG dominates as expected and to flag annotations in which non-canonical starts are unusually frequent before drawing conclusions from the downstream metrics.

### 2.3 Motif construction and positional statistics

From the extracted start codon contexts, position-specific nucleotide frequencies *P_i_*(*b*) (with *b* ∈ {*A, C, G, T* } and *i* a position in the analysis window) are computed and assembled into the PFM. A sequence logo is rendered from the PFM using the `logomaker` Python library [21]; the per-position letter heights follow the information-content convention described below in the Information content subsection.

In addition to the nucleotide-level logo, the application produces complementary purine/pyrimidine aggregated profiles to highlight broader compositional trends. These aggregated profiles collapse the four nucleotides into two purine bases (A, G) and two pyrimidine bases (C, T), revealing coarser-grained compositional biases that can be masked by the full four-letter alphabet. Figure 1 illustrates this complementary view for *Saccharomyces cerevisiae*.

**Figure 1:**
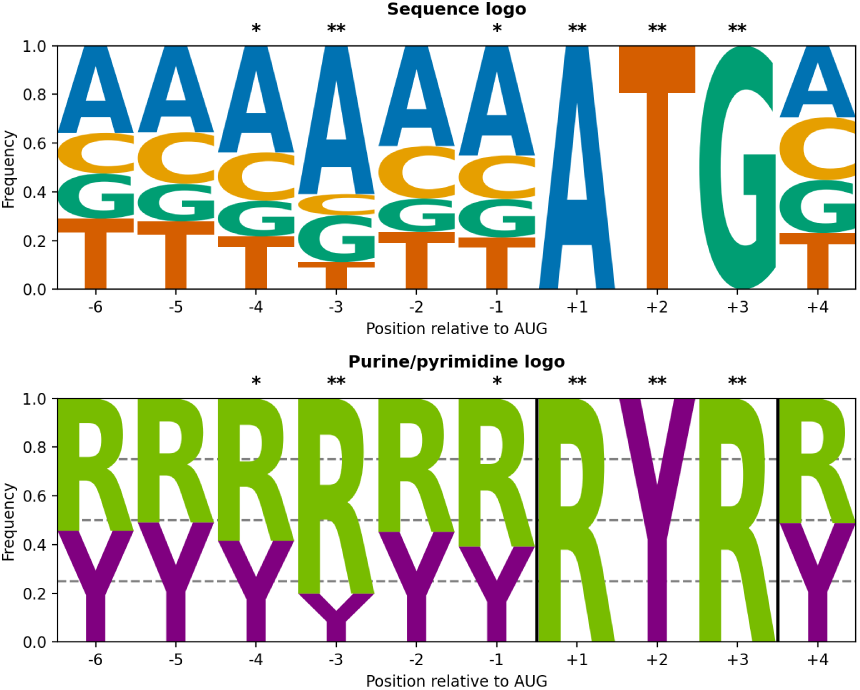
Complementary visualization of translation initiation context in *Saccharomyces cerevisiae* across positions −6 to +4 relative to the AUG start codon. Top: standard sequence logo showing per-position nucleotide frequencies. Bottom: purine/pyrimidine aggregated logo collapsing the four nucleotides into two purine bases (A, G) and two pyrimidine bases (C, T) to highlight broader compositional trends. Both panels are generated using the sequana. `kozak` module (methods `plot_logo and plot_logo_purine_pyrimidine`). Stars above each position indicate chi-square significance against the genome-wide nucleotide background (∗: *p <* 0.05; ∗∗: *p <* 0.01). The canonical Kozak position at −3 is visible in both representations.

To assess positional deviations from background composition, a chi-square test is applied at each position using the genome-wide nucleotide frequencies of the selected genome as the null model. The resulting statistics provide an exploratory measure of enrichment relative to that genome’s baseline composition. Significance markers (∗: *p <* 0.05; ∗∗: *p <* 0.01) are overlaid above each position (Figure 1) to highlight positions whose nucleotide composition deviates significantly from the genomic background. No correction for multiple comparisons is applied; the markers are intended as exploratory guides rather than hypothesis-testing outcomes.

### 2.4 Kullback–Leibler divergence and the Kozak Strength Index

Positional nucleotide constraints are quantified using Shannon entropy and KL divergence, both computed from the PFM of the selected genome.

Entropy at position *i* is

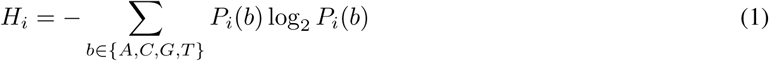

where *P_i_*(*b*) represents the nucleotide frequency at position *i*.

To account for the nucleotide bias of the selected genome, KL divergence is computed relative to a background distribution:

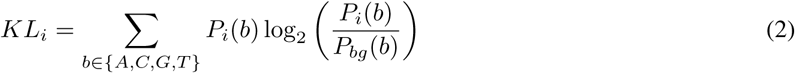

where *P_bg_*(*b*) denotes the background frequency. Three background models are exposed in the interface: (i) the genomic nucleotide background, (ii) an in-frame ATG background built from internal ATG codons of the same genome (the annotated start sites are removed), and (iii) the same in-frame ATG pool with its sequences shuffled. The user picks the model best suited to the genome under inspection. Model (ii) is in principle the most natural null because its constituent windows already contain an ATG in the same reading frame, so its base composition is closer to the local context of a true start codon than the genome-wide average. However, for GC-rich bacterial genomes in particular, random in-frame ATG contexts inherit codon-position biases of the host genome and produce a characteristic 3-bp periodicity which contaminates the KL profile [9, 13]; in that situation, the genome-wide background gives a more stable baseline. To address this trade-off, model (iii) keeps the same in-frame ATG pool but shuffles each sequence per row, which destroys the codon-frame periodicity while preserving the per-window composition, recovering the true localized signal rather than the 3-bp artifact. An illustration on *S. cerevisiae* is shown in Figure S1 (Supplementary Information): the canonical −3 peak is recovered with all three backgrounds, but its amplitude varies depending on the null, demonstrating that the choice of background changes the magnitude—not the location—of the inferred constraint; the same comparison on a GC-rich bacterial genome, where the artifact is most severe, is shown in Figure S2 (Supplementary Information).

For genomes annotated with few coding sequences, bootstrap resampling can be enabled to estimate uncertainty on each *KL_i_* value. For genomes annotated with thousands of genes, the estimates are stable and bootstrap is disabled by default to keep the interactive response time low.

From the per-position *KL_i_*profile, several scalar summaries characterize the intensity, localization, and spatial organization of the initiation signal of the selected genome. The headline summary is the **Kozak Strength Index (KSI)**, defined as the mean KL divergence across the six nucleotides immediately upstream of the start codon, [imline1] KSI captures localized constraints around the start codon: high values correspond to strong, focused motifs typical of eukaryotic Kozak sequences. The KSI window (−6 to −1) spans the six positions of the canonical Kozak context; positions farther upstream (−20 to −7) and downstream (+4 to +6) are retained in the full feature vector for the embeddings but excluded from KSI to maintain focus on the eukaryotic mechanistic core. Conceptually related per-sequence scoring schemes have been proposed, such as the Kozak Similarity Score (KSS) of [22], which ranks individual transcripts and was developed to identify alternative translation initiation codons implicated in cancer. The KSS is computed on a per-sequence basis using a position-weighted consensus matrix *W_i_*(*b*):

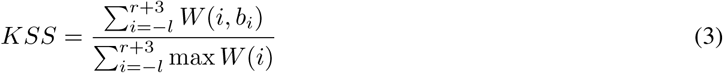

where the index *i* runs over the *l* + 3 + *r* positions of the analysis window centered on the start codon (upstream positions −*l* to −1, the three ATG positions, and downstream positions +4 to *r* + 3). *b_i_*is the nucleotide at position *i* in a given sequence, and *W* (*i, b*) is the position-specific weight assigned to nucleotide *b* at position *i*, read directly from the matrix published by [22]. *W* (*i*) denotes the four-component vector (*W* (*i, A*)*, W* (*i, C*)*, W* (*i, G*)*, W* (*i, T* )) at position *i*, and max *W* (*i*) is its largest entry. *W* (*i, b*) is therefore an externally provided reference matrix, not a quantity estimated from our corpus. The reference matrix carries zero weight at the three ATG rows, so those positions contribute neither to the numerator nor to the denominator and the score is effectively driven by the flanking nucleotides. The denominator normalizes the score so that KSS values fall in [0, 1], where higher scores indicate closer agreement with the strong Kozak consensus. The implementation in `sequana.kozak` uses *l* = *r* = 10 by default (a 23-base window matching the original [22] matrix); these flanks are configurable on the `KozakWeightScore` class.

KSI differs from KSS in that KSI is a genome-level information-theoretic summary reflecting observed constraints across all annotated start codons of the genome under analysis, whereas KSS is a per-transcript score that uses the externally derived weight matrix to rank individual sequences.

Auxiliary scalar summaries derived from the same KL profile—total upstream information *I_total_*, peak information *I_peak_*, signal concentration *C* = *I_peak_/I_total_*, and effective half-information width *W*_50_—are defined in the Supplementary Information (Auxiliary genome-level scalar metrics). They are computed and exported by the software for downstream comparison but are not used in the main-text figures or in the t-SNE feature vector.

### 2.5 Information content (IC) per position

In addition to entropy and KL divergence, the application reports the IC at each position, a measure of sequence conservation in bits:

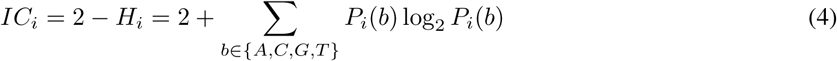

where *H_i_*is the Shannon entropy at position *i*. IC ranges from 0 bits (uniform nucleotide distribution) to 2 bits (complete conservation to a single nucleotide). One could equivalently report *H_i_* alone, but information content is preferred here for three reasons: (i) it is monotonically aligned with constraint—larger *IC_i_* means a more constrained position, whereas larger *H_i_* means a less constrained one, which is counter-intuitive when comparing peaks across genomes; (ii) IC has the same units (bits) as the KL divergence used elsewhere in the analysis, so the two profiles can be plotted on a common axis and compared directly; (iii) IC is the standard letter-height convention for sequence logos [23], and using the same quantity in the per-position profile and in the rendered logo keeps the numerical summaries and the visual representation consistent. The same *IC_i_* values drive the sequence logo: each nucleotide letter at position *i* is rendered with height *P_i_*(*b*) · *IC_i_*. From the IC profile of the selected genome, three summary scalars are derived:

- **Maximum IC (***IC_max_*): The highest IC value across the upstream window, indicating the strongest positional constraint.
- **Total IC (***IC_sum_*): The sum of IC values across all positions, capturing cumulative sequence constraint.
- **Position of maximum IC**: The upstream position at which the strongest constraint occurs.

Although KSI, the IC summaries, and the auxiliary KL-derived metrics (Supplementary Information) are all derived from the same PFM, they are not redundant: they capture different facets of the initiation signal (overall intensity, peak height, localization, width, positional bias). A Spearman correlation analysis of these metrics across the 2,282 eukaryotic reference genomes (Figure S3) shows the expected strong coupling between KSI and total upstream information (*I_total_*), and between *IC_max_*and *I_peak_*, but reveals that signal concentration (*C*), peak position, and GC content carry largely independent information. This justifies retaining the full panel of metrics for downstream comparative analysis rather than collapsing them to a single summary.

### 2.6 Context-strength classification

As an interpretable, position-anchored complement to the continuous metrics above, KozakExplorer also assigns each genome’s consensus context a qualitative tier following the strong/adequate/weak scheme, widely adopted following Kozak’s original analysis [3, 6]. The classification rests on the two functionally dominant positions of the eukaryotic Kozak context—a purine (*R*) at −3 and a *G* at +4 relative to the AUG: a context is labelled *strong* when both are present, *adequate* when exactly one is, and *weak* when neither is. Strong contexts that additionally match the full GCCRCC upstream vertebrate consensus [3] are further labelled *optimal*. The tier is computed by the `classify_kozak_strength` routine of `sequana.kozak` and, aggregated across a corpus, yields the fraction of genomes in each tier, summarizing how closely the species-level consensus approaches the canonical eukaryotic hallmark.

### 2.7 Precomputed multi-genome corpus and landscapes

Beyond single-genome interactive analysis, KozakExplorer ships with precomputed metric tables and t-SNE embeddings for a large reference corpus. Reference genomes were retrieved from the NCBI Genome database using the RefSeq reference genome filter. Only assemblies designated as reference genomes and accompanied by curated Ref-Seq annotations were included. Accessions were obtained programmatically using the NCBI Datasets interface with the filters `reference_only=true, annotated_only=true, and refseq_annotation=true`. This strategy ensured standardized, high-quality annotations suitable for large-scale comparative analysis. Three landscapes are distributed:

#### Primary (eukaryota)

2,282 eukaryotic reference genomes constitute the primary focus of this study, enabling detailed investigation of Kozak landscape patterns across kingdoms.

#### Full (eukaryota + bacteria + archaea)

An additional 22,253 bacterial and 809 archaeal reference genomes were included to provide comparative context and reveal global transitions in translation initiation architecture across domains of life (25,344 genomes total); a per-kingdom breakdown of this full corpus is provided in Table S1 (Supplementary Information).

#### Apicomplexa case study

A dedicated dataset of 216 Apicomplexa genomes was independently retrieved from NCBI by combining RefSeq and GenBank assemblies under looser filtering criteria than the main eukaryotic dataset, to maximize representation of this clinically important parasite-rich clade. This separate pull provides broader Apicomplexa coverage than filtering the main 25,344-genome dataset by phylum. The genus/subgenus classification rules are summarized in Table S2, and the list of all 216 assemblies and their basic metadata is provided in the GitHub repository https://github.com/sequana/webapp_kozak.

All accession lists (eukaryota, full cross-domain, and Apicomplexa) are distributed as plain-text files in the `webapp_kozak` repository (https://github.com/sequana/webapp_kozak) and were used as the input for the automated analysis pipeline described next.

Taxonomic classifications and assembly statistics (full taxonomic lineage, GC content, number of contigs, total assembly length, contig and scaffold N50 values) were retrieved from NCBI metadata resources and integrated with Kozak-derived metrics to enable meta-analysis of the comparative “Kozak landscape”.

### 2.8 Automated analysis workflow

The three landscapes were generated using a reproducible workflow [24, 25] implemented in Snakemake. The Snakefile and per-rule scripts, together with the accession lists consumed as input, are tracked alongside the ap-plication in the `webapp_kozak` repository (https://github.com/sequana/webapp_kozak; see in particular `models/Snakefile`), so the full corpus regeneration can be reproduced from the published sources. The pipeline automates genome retrieval, extraction of translation initiation contexts, computation of statistical metrics, and aggregation of results. To keep all genomes commensurate, the pipeline fixes the extraction window to [−20, +6] around the start codon (whereas the interactive web application leaves these bounds to the user). The choice of [−20, +6] is arbitrary but pragmatic: the upstream span comfortably covers both the canonical eukaryotic Kozak determinants centered near −3 [2, 3, 4] and the bacterial and archaeal Shine–Dalgarno region typically located 5–13 nucleotides upstream [10, 13], while the downstream span captures the +4 position known to modulate initiation efficiency [3]. Only canonical ATG starts are retained, again to keep cross-genome comparisons uniform.

For each genome, the workflow:

1. downloads genome FASTA and GFF3 annotation files,
2. extracts annotated start codon contexts,
3. computes positional nucleotide frequencies,
4. calculates information-theoretic metrics,
5. generates motif visualizations and statistical summaries,
6. merges genome-level results into a global comparative dataset.

All analyses were performed through the kozak module of the Sequana library, so single-genome interactive results and batch corpus results are computed identically.

### 2.9 Performance and scalability

The pipeline was executed on an institutional SLURM cluster (Snakemake 9.19, up to 200 concurrent jobs). Processing the full reference set of 25,344 genomes required approximately 234 CPU-days of aggregate compute time (5,610 CPU-hours). NCBI metadata and asset retrieval dominated wall-time consumption (∼60% of CPU-hours, throttled by remote API rate limits), while the per-genome scientific computation accounted for the remaining ∼40% (∼2,219 CPU-hours, median ∼1.8 min per genome). Per-genome memory usage was modest (median peak RSS 293 MB) but exhibited a long tail driven by large vertebrate and polyploid plant assemblies, reaching ∼51 GB on the largest genomes. This ∼51 GB figure is the peak RSS of the single largest assembly processed; the vast majority of genomes complete in under 1 GB, and large-RAM nodes were required only for a handful of outlier assemblies. Such genomes can equally be processed on a commodity workstation by running them individually rather than concurrently. The full output collection occupies 12 GB of intermediate artifacts; the consolidated metric and embedding tables distributed with the publication compress to approximately 52 MB across four Parquet files. The pipeline supports both local and SLURM execution modes; per-rule resource declarations are provided so that mid-range workstations can reproduce the workflow on smaller subsets without modification.

### 2.10 Comparative analysis and dimensionality reduction

To explore patterns across eukaryotic genomes and comparative contexts, genome-level feature vectors were standardized and embedded using Principal Component Analysis (PCA), Uniform Manifold Approximation and Projection (UMAP), and t-distributed Stochastic Neighbor Embedding (t-SNE). PCA is deterministic—given the same feature matrix it always returns the same components—and serves as a useful reproducibility baseline; however, on our genome-level feature vectors the first two principal components explain only a modest fraction of the variance and the kingdom-level groups overlap heavily, so PCA fails to resolve the cluster structure that we want to expose. Two non-linear alternatives, UMAP and t-SNE, drop the linearity assumption and preserve local neighborhood structure—the relevant regime here, since the mapping from feature differences to lineage differences is non-linear. We retained t-SNE as the headline embedding for two practical reasons. First, t-SNE’s local structure is governed by an explicit, tunable perplexity parameter, which gave more reproducible cluster boundaries on this dataset than UMAP’s default *n*_*neighbors*/*min*_*dist* trade-off. Second, both UMAP and t-SNE are stochastic, but t-SNE’s variability is well characterized and easy to tame with a consensus procedure (described below). UMAP and PCA outputs are still computed and stored for cross-checking. Three separate embeddings were produced with this protocol: (1) eukaryota-only, fit on 2,282 eukaryotic genomes to highlight eukaryotic-specific structure; (2) full, fit on all 25,344 genomes for global comparative context; and (3) apicomplexa-dedicated, fit on 216 Apicomplexa genomes for focused parasite analysis.

The same per-genome feature vector is used as input for all three embeddings (PCA, UMAP, t-SNE): the concatenation of three per-position blocks evaluated on the analysis window: (i) the KL divergence profile {*KL_i_*}, (ii) the informa-tion content profile {*IC_i_*}, and (iii) the position-specific motif matrix {*P_i_*(*b*)}*_b_*_∈{_*_A,C,G,T_* _}_. For the default window (−20, +6, with the AUG itself excluded by `include_start_codon=False`), each profile spans 26 positions, so the final feature vector concatenates 6 per-position vectors (one for KL, one for IC, and four for the motif nucleotide frequencies) for a total of 156 features. The three blocks are standardized independently before concatenation so that no single block dominates the pairwise distances used by the embedding. Importantly, the genome-level summary metrics defined above (KSI, *IC_max_*, *IC_sum_*, peak position, GC content) and the auxiliary metrics in the Supplementary Information (*I_total_*, *I_peak_*, *C*, *W*_50_) are *not* part of the embedding feature vector; they are computed from the same KL and IC profiles and used downstream as coloring overlays and per-genome scalar comparisons. To address t-SNE stochasticity in the smaller-clade analysis, we employed consensus t-SNE: running 60 independent t-SNE seeds at perplexity 15, aligning each output to a reference via Procrustes transformation (centering, scaling, orthogonal rotation), and averaging aligned coordinates. This approach yields stable consensus coordinates and per-point uncertainty (standard deviation across runs) for identifying unreliable embeddings. The consensus procedure is used for the Apicomplexa-specific embedding, where the dataset is small enough (216 genomes) that single-run variability is non-negligible. For the eukaryota-only and full cross-domain landscapes, the larger sample sizes (2,282 and 25,344 genomes respectively) make the embedding much more stable; these were computed with the kozak build_embeddings command of webapp_kozak at perplexity 50, and the figures show a single representative run rather than a Procrustes-averaged consensus.

## 3 Results

### 3.1 KozakExplorer: interactive page and precomputed landscapes

The KozakExplorer web application offers two complementary entry points. **The landing page** lets a user analyze a single genome on demand (e.g. a local genome and annotation or a RefSeq accession); the application extracts the metrics and renders the corresponding plots interactively (see Materials and Methods). This single-genome capability is itself a primary result of this work: from nothing more than a RefSeq accession or a local FASTA/GFF3 pair, KozakExplorer returns not only the conventional sequence-logo motif but also the per-position KL divergence and information-content profiles that quantify the strength and localization of the initiation signal. Figure 2 illustrates this output for three contrasting organisms—*Leishmania major* (weak, dispersed signal), *Saccharomyces cerevisiae* (strong, focused eukaryotic Kozak), and *Escherichia coli* (distributed Shine–Dalgarno pattern)—each rendered as the logo, KL profile, and IC profile. **The companion landscape pages**, by contrast, expose precomputed metric tables and t-SNE embeddings over large reference corpora—2,282 eukaryotic, 25,344 cross-domain, and 216 Apicomplexa genomes—produced offline by the Snakemake pipeline described in Materials and Methods (Automated analysis workflow subsection). All Results subsections below draw on those precomputed landscapes.

**Figure 2:**
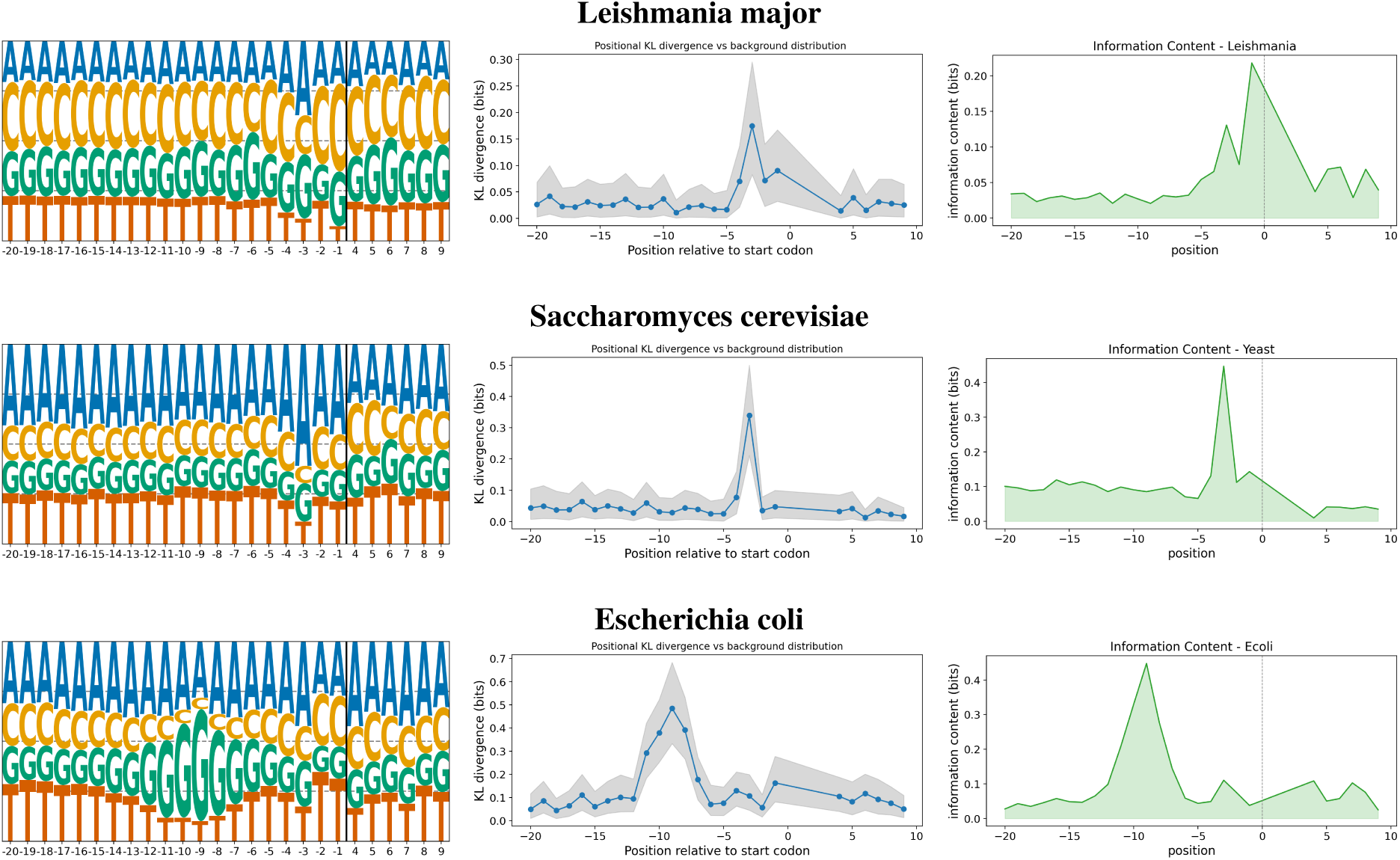
Comparative analysis of translation initiation contexts using KozakExplorer. Left panel: sequence logo of per-position nucleotide frequencies. Center panel: KL divergence profile quantifying positional information relative to the genomic background. Right panel: information content per position, in bits. Top row: *Leishmania major* (parasitic protist, eukaryotic Kozak, weak signal). Middle row: *Saccharomyces cerevisiae* (yeast, canonical eukaryotic Kozak, strong focused motif). Bottom row: *Escherichia coli* (bacterium, Shine–Dalgarno signal, distributed upstream pattern).

### 3.2 Per-genome initiation signals: strength, position, and diversity

Among eukaryotic genomes, the KSI exhibited substantial variability, reflecting diversity in selective pressure on the KCS and translation initiation context (Figure 5, bottom-right panel). Peak position, however, remained predominantly conserved at nucleotide position –3 relative to the AUG start codon across phylogenetically distant eukaryotic lineages, as illustrated both for individual genomes (*Leishmania major* and *Saccharomyces cerevisiae*; Figure 2) and across kingdoms (Figure 3; Table 1).

**Figure 3:**
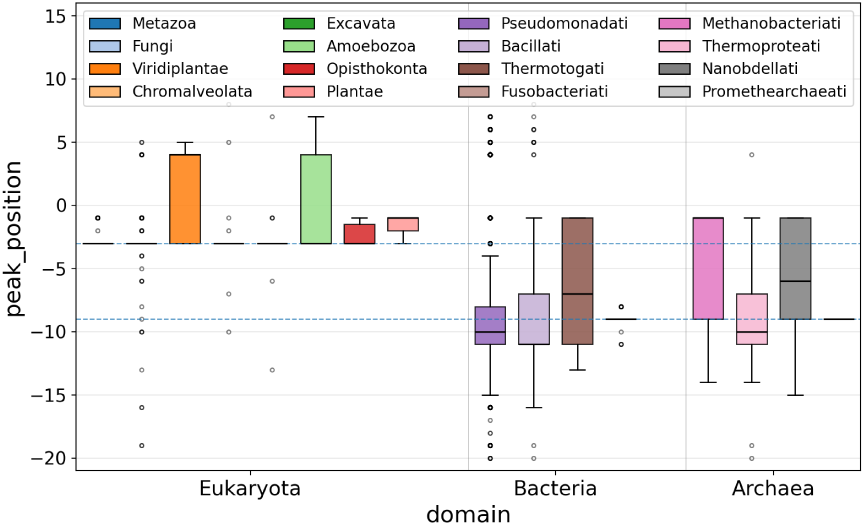
Distribution of peak position across analyzed genomes. The wide dispersion illustrates the diversity of ini-tiation signal positions observed across taxa. Eukaryotic genomes generally display stronger and localized constraints at the canonical position −3, whereas most bacterial and archaeal genomes show broader positional patterns consistent with Shine–Dalgarno sequences.

**Figure 4:**
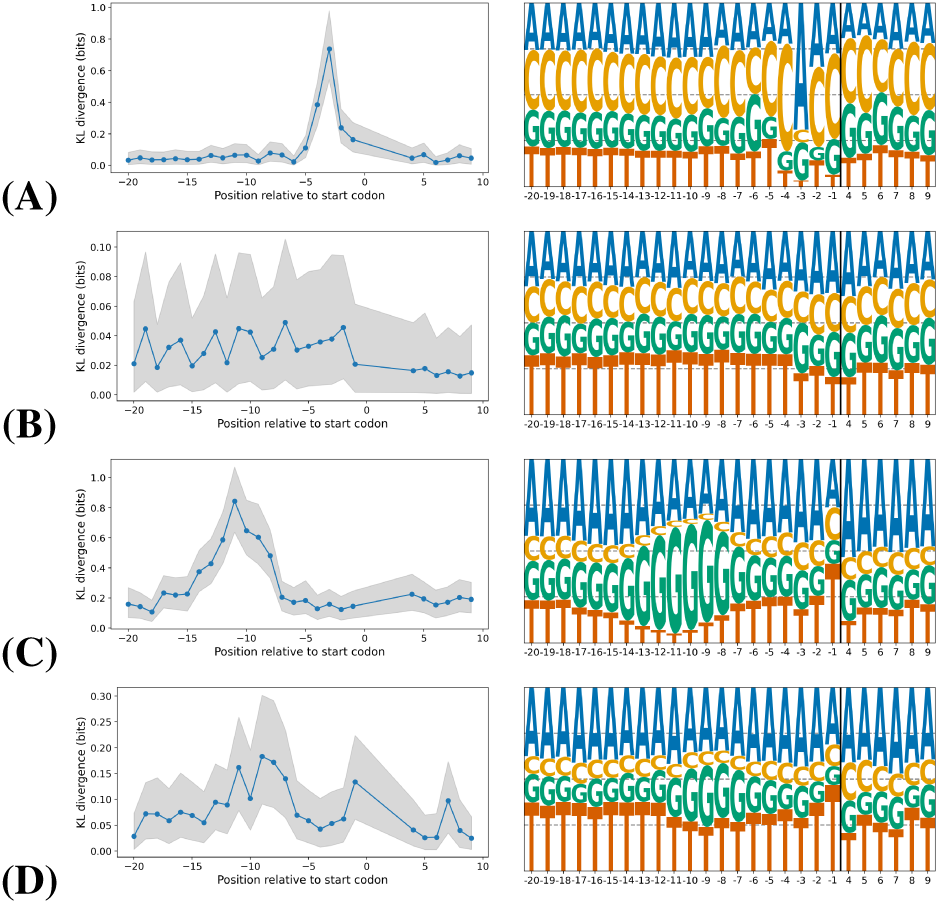
Representative positional information landscapes and sequence logos illustrating the diversity of initiation signal architectures. Each row shows one genome with its KL divergence profile (left) and sequence logo (right). From top to bottom: (A) a eukaryotic genome with a strong Kozak-like peak, (B) a eukaryotic genome with weaker positional constraints, (C) a bacterial genome with a narrow upstream signal, and (D) a bacterial genome with a broader or weakly defined initiation context.

**Figure 5:**
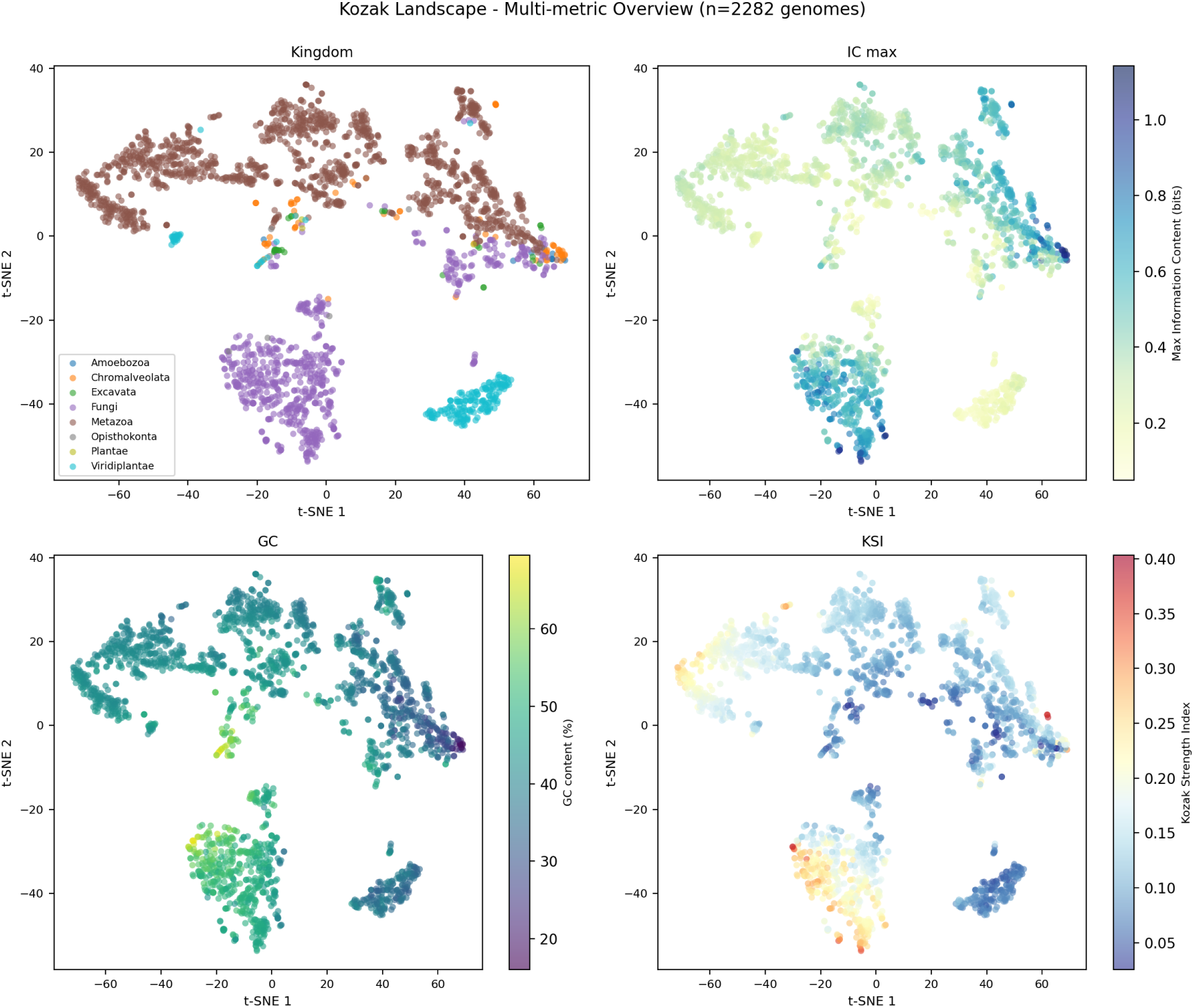
t-SNE embedding (single representative run, perplexity 50, computed with kozak build_embeddings) of 2,282 eukaryotic reference genomes based on the per-genome concatenation of three per-position vectors: KL diver-gence profile, information content profile, and motif nucleotide frequencies *P_i_*(*b*). Genome-level summary metrics (KSI, *IC_max_*, GC content, kingdom) are used only to color the embedding and are not part of the feature vector. Top-left: Kingdom-level clustering (subset of kingdoms shown: Metazoa, Fungi, Viridiplantae, Chromalveolata, Excavata, Amoebozoa). Top-right: Maximum information content (showing positional sequence conservation). Bottom-left: GC content (showing genomic composition effects). Bottom-right: Kozak Strength Index (showing kingdom-level variation in signal strength). Continuous metrics use viridis-family colormaps; categorical variables use distinct colors per group. All genomes project into a structured landscape revealing both kingdom-level organization and continuous variation in translation initiation architecture.

**Table 1:**
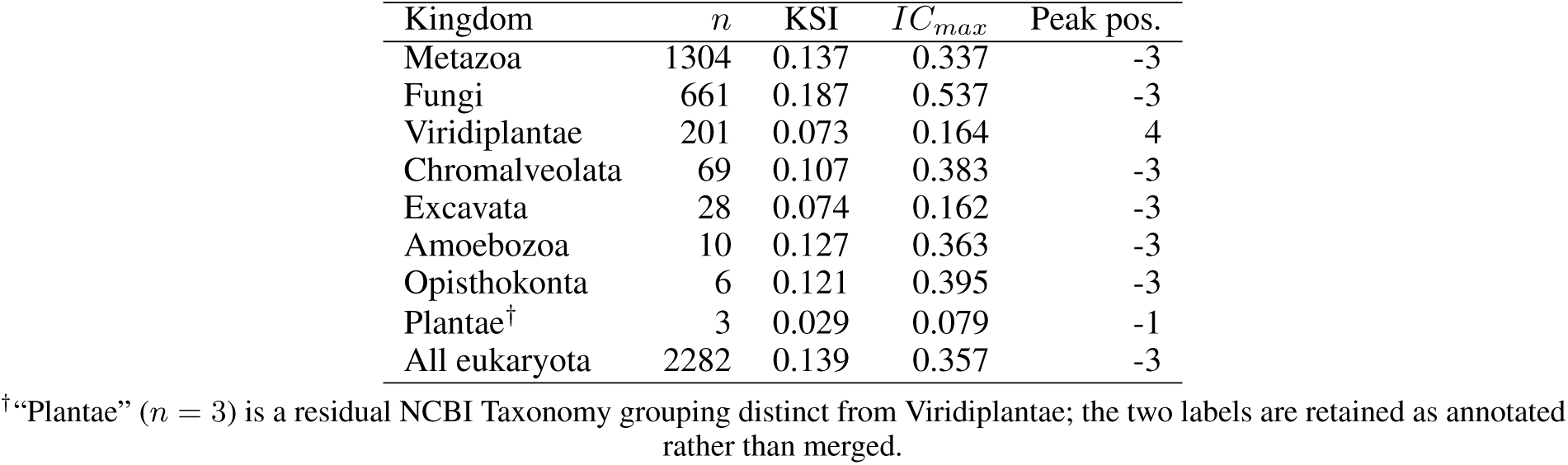
Per-kingdom summary of translation initiation metrics across the eukaryotic dataset. KSI: Kozak Strength Index (median across genomes); *IC_max_*: median of the maximum per-position information content, in bits; Peak pos.: mode of the peak position across genomes, in nucleotides relative to the AUG start codon. The KSI and *IC_max_* columns therefore report per-kingdom median values; the “median” qualifier is dropped from the header for compactness. Interquartile ranges of KSI are reported in Table S1 (Supplementary Information).

IC-derived metrics (*IC_max_*, *IC_sum_*) similarly varied widely across eukaryotic genomes, highlighting diversity in the strength and spatial organization of Kozak constraints. Table 1 reports per-kingdom summary statistics: Fungi exhibit the strongest signals (median KSI = 0.187, *IC_max_* = 0.537 bits) consistent with the canonical yeast Kozak motif, whereas Viridiplantae and Excavata display markedly weaker constraints (median KSI ≈ 0.07). The peak position is conserved at −3 in all kingdoms with *n >* 3 except Viridiplantae, where the modal peak shifts to +4. This is not an artifact: position +4 is independently known to carry a strong eukaryotic KCS bias (typically a *G* [3, 4]), and in plant genomes the upstream −3 signal is weak enough (median KSI ≈ 0.07) that the residual downstream +4 constraint becomes the dominant per-position peak.

KL divergence profiles revealed distinct architectural patterns of translation initiation signals. Some genomes exhibited sharply localized peaks immediately upstream of the start codon, characteristic of strong KCS-like constraints. Others displayed broader upstream enrichment patterns consistent with distributed regulatory signals.

Representative examples illustrating these contrasting profiles are shown in Figure 4. For instance, some eukaryotic genomes display strong and narrow upstream information peaks, whereas others exhibit weaker or more diffuse constraints (panels A–B). Among bacterial genomes, both sharply localized and broadly distributed signals can be observed, and in some cases the positional signal is minimal (panels C–D).

### 3.3 Eukaryotic t-SNE embedding and kingdom-level organization

To explore the diversity of translation initiation signals within Eukaryota, we projected genome-level metrics into a low-dimensional space via t-SNE (Figure 5). This embedding, fit exclusively on eukaryotic genomes, reveals clear kingdom-level clustering (Metazoa, Fungi, Viridiplantae, Chromalveolata, Excavata, Amoebozoa, Opisthokonta, Plan-tae) and continuous variability in signal strength and organization among eukaryotes.

Information content (IC) varies across kingdoms, reflecting lineage-specific adaptations in translational constraint stringency. The integration of IC metrics into the t-SNE embedding distinguishes eukaryotic genomes with highly conserved motifs (high IC*_max_*, concentrated information) from those with more diffuse constraints (lower IC*_max_*, distributed information).

The embedding is shown as four panels, each coloring the same projection by a different genome-level variable. The top-left panel (kingdom) reveals fine-grained structure within kingdoms: notably, within Viridiplantae (green plants), the Streptophyta phylum exhibits a striking bipartition into two spatially distinct regions. This pattern corresponds to the evolutionary split between monocots (grasses, lilies) and dicots (legumes, eudicots), suggesting that Kozak landscape divergence mirrors major transitions in plant body plans and gene expression architectures (quantified in the Consistency subsection). The top-right panel (IC*_max_*) highlights positional conservation: Fungi and some Metazoa cluster with high information content, indicating highly constrained motifs, while other eukaryotic lineages exhibit more diffuse information distributions. The bottom-left panel (GC content) shows compositional effects on the landscape, reflecting the broad range of genomic GC

### 3.4 Consistency with known Kozak biology

To validate that the KSI captures biologically meaningful patterns, we examined the distribution of KSI across eukaryotic kingdoms (Table 1). The results align with established understanding of translation initiation: Fungi exhibit the strongest median KSI (0.187, IQR 0.125–0.225), reflecting the choice of the [−6, −1] window motivated by canonical yeast Kozak literature; this outcome aligns with known selective pressure on eukaryotic translation initiation in model fungi, validating that KSI captures biologically expected patterns. Metazoa show the second-highest median KSI (0.137, IQR 0.109–0.170), reflecting strong selective pressure on eukaryotic translation initiation (per-kingdom IQRs are tabulated in Table S1, Supplementary Information). In contrast, Excavata and Viridiplantae display markedly weaker KSI signals (medians 0.074 and 0.073, respectively). This matches prior observations that plants employ more degenerate or variant initiation contexts [26, 27], while kinetoplastids rely on post-transcriptional rather than canonical scanning-based regulation [28, 29, 30].

The peak position remains structurally conserved at −3 across all kingdoms, confirming that recognition of the −3 purine context is a universal feature of scanning-dependent eukaryotic translation initiation. This invariance across otherwise divergent KSI values underscores the distinction between the positional architecture (hard structural constraint) and signal strength (tunable evolutionary parameter).

At the genome level, extreme examples further validate the metric. The highest-KSI genomes include *Naegleria lovaniensis* (Excavata, KSI = 0.403) and multiple Fungi species (KSI ≥ 0.34), while lowest-KSI genomes are dominated by Amoebozoa (*Entamoeba* spp., KSI ≈ 0.025). The correlation between KSI and maximum information content (*IC_max_*, Pearson *r* = 0.598) indicates that KSI captures genuine sequence constraint rather than noise, and validates its use as a genome-wide summary statistic suitable for comparative analysis.

A particularly stringent validation is provided by the monocot–dicot split within Streptophyta, the most striking sub-kingdom signal recovered without supervision in this study: the eukaryotic embedding (Figure 5) divides Viridiplantae into two non-overlapping regions corresponding to monocots and dicots. This recovers, at substantially larger sample size, contextual differences previously reported for plant translation initiation [31, 7]. Our corpus comprises 150 dicot and 33 monocot genomes—well beyond the 7 monocot and 7 dicot species analyzed by Gupta et al. [31]—and the per-position consensus both confirms and sharpens their observations. At position −3, all 150 dicots carry an A, whereas 85% of monocots carry a G (the remaining 15% an A), recovering the reported −3A (dicot) versus −3G (monocot) bias. The +4G is universal in both clades; every dicot reaches the *strong* context tier, while monocots split evenly between the *strong* and *optimal* tiers, the latter reflecting their GC-rich upstream context matching the full GCCRCC consensus. The −2 position is conserved as a M (A or C) in both clades, as noted by Hernández et al. [7], but flips between them—predominantly A in dicots and C in monocots—and position −4 is uniformly A in dicots while monocots again split between C and A, consistent with their GC-shifted composition. The split is unlikely to be a pure GC-content artifact, since the GC-colored panel in Figure 5 (bottom-left) does not reproduce the bipartition cleanly.

### 3.5 Context-strength tiers across kingdoms

Classifying each genome’s consensus context into the optimal/strong/adequate/weak tiers (Section 2.6) summarizes how closely a species-level consensus approaches the canonical eukaryotic hallmark. Across the 2,282 eukaryotic genomes, nearly half (1,125; 49.3%) reach the optimal or strong tier—carrying both the −3 purine and the +4 G—while weak contexts are rare (15 genomes, 0.7%), confirming that the hallmark is widespread across eukaryotic diversity. Resolving the corpus by kingdom (Table 2), however, shows that the tier classification and the KSI capture complementary, and sometimes opposing, aspects of the initiation context. Metazoa follow the expected pattern, with most genomes (908 of 1,304) reaching the strong or optimal tier, consistent with their high KSI. The two measures diverge sharply for Fungi and Viridiplantae. Fungi, despite carrying the strongest upstream signal in the corpus (median KSI 0.187; Table 1), contain *no* strong or optimal genomes: their consensus satisfies the −3 purine but lacks the +4 G that the tier requires, so all 661 genomes fall in the adequate or weak tiers. Viridiplantae show the opposite inversion—a weak upstream signal (median KSI ≈ 0.07) yet 194 of 201 genomes in the strong or optimal tier, because both anchor positions (−3 purine and +4 G) are satisfied even though the surrounding −6 to −1 window is poorly constrained; this is the same +4 G that shifts the plant per-position peak to +4 (Table 1). The two descriptors are therefore not redundant: KSI summarizes the breadth of the upstream −6 to −1 constraint, whereas the tier is anchored on the two positions of greatest mechanistic importance—including the downstream +4, which lies outside the KSI window.

**Table 2:**
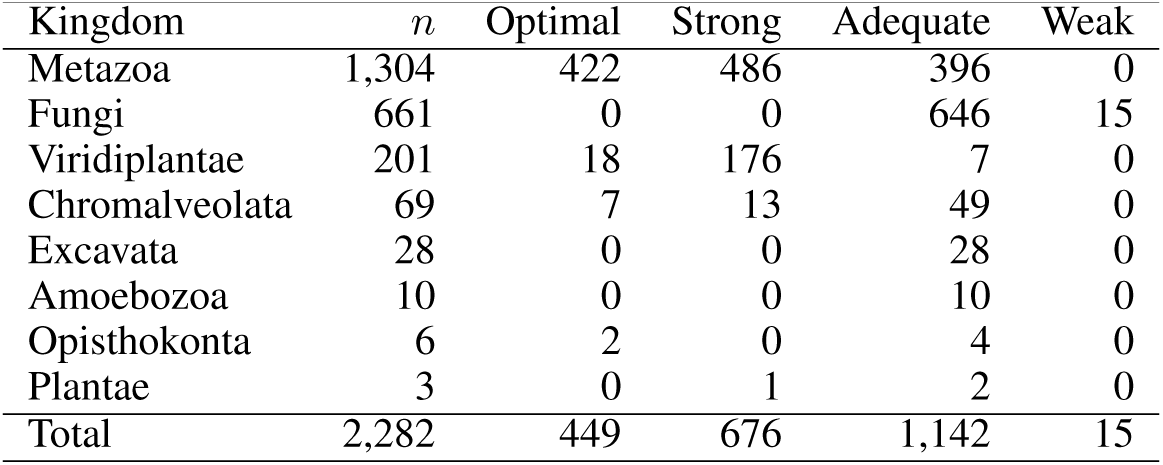
Per-kingdom distribution of Kozak context-strength tiers across the 2,282 eukaryotic reference genomes. Counts are numbers of genomes assigned to each tier (optimal, strong, adequate, weak) following the strong/adequate/weak scheme [6] (Materials and Methods); *n* is the number of genomes in the kingdom. Note the contrast between Fungi (no strong/optimal genomes despite the highest median KSI in Table 1) and Viridiplantae (mostly strong/optimal despite a weak median KSI).

The dominance of the −3 purine that underlies the strong and adequate tiers also lets us address a question raised as an outstanding issue by [7]: is the purine at the −3 position conserved across all eukaryotes? Across the 2,282 eukaryotic genomes, a purine (A or G) is the majority base at −3 in 99.2% of genomes (2,264 of 2,282; median purine fraction 81%, interquartile range 78–84%), and the bias is present in every kingdom—including Viridiplantae (mean 70%), whose upstream window is otherwise weakly constrained. The conservation is therefore near-universal but not absolute: the 18 genomes (0.8%) in which the −3 purine falls below 50% are almost exclusively Microsporidia (17 genomes), together with a single parabasalid (*Histomonas meleagridis*)—all highly reduced, AT-rich intracellular parasites with divergent translation machinery. The −3 purine thus behaves as a near-invariant anchor of the eukaryotic initiation context that erodes only in the most derived parasitic lineages.

### 3.6 Apicomplexa: a case study of parasitic protist translation initiation

Apicomplexa (*Plasmodium*, *Toxoplasma*, *Cryptosporidium*, *Babesia* and *Theileria*) represent a clinically important eukaryotic clade with strong host specialization and known divergence in gene expression mechanisms. We performed a dedicated analysis of 216 Apicomplexa genomes retrieved from NCBI (combining RefSeq and GenBank assemblies) using looser filtering criteria than the main eukaryotic dataset to maximize clade representation. This Apicomplexa-specific embedding (Figure 6) reveals distinct clustering structured by genus, in line with host range and phylogeny.

**Figure 6:**
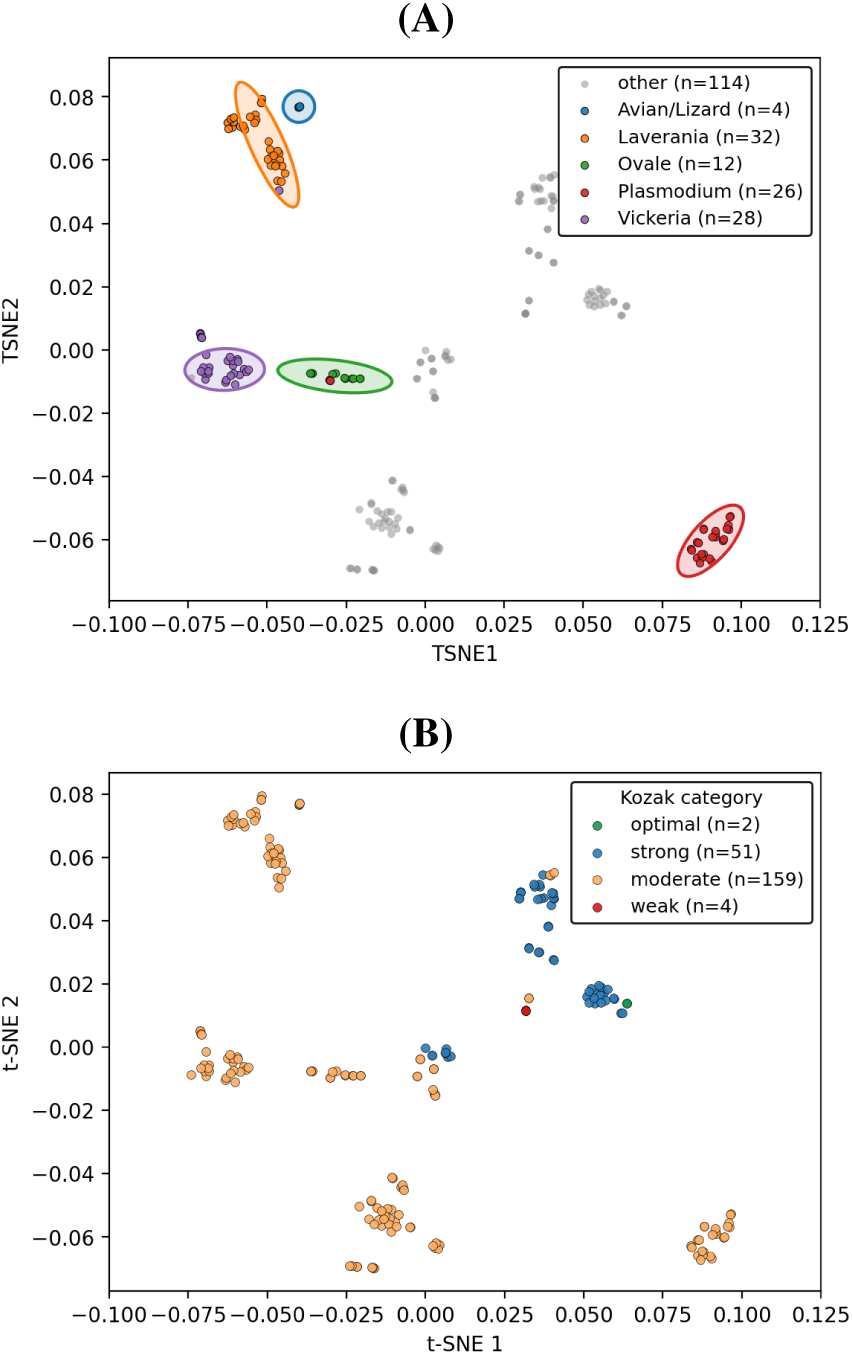
t-SNE embedding of 216 Apicomplexa genomes (NCBI RefSeq + GenBank) over a single set of 60-run consensus coordinates, shown with two colorings. **(A)** Colored by genus and, for *Plasmodium*, by subgenus (Laverania, Vinckeia, Ovale, primate-infecting clade). Genus- and subgenus-level structure is recovered, reflecting host range and phylogeny. The single gray point inside the Vinckeia cluster is a *Hepatocystis* genome, expected to lie close to the rodent-malaria *Plasmodium* subgenus given its known phylogenetic placement. Subgenus assignments follow the rules in `apicomplexa_classification.json` (see https://github.com/sequana/webapp_kozak). **(B)** The same embedding colored by Kozak context-strength tier (optimal/strong/moderate/weak; Section 2.6); the strong-tier genomes form a coherent cluster, showing that context strength is spatially structured in the embedding rather than randomly distributed.

Within *Plasmodium*, the embedding separates the major subgenera (Laverania, Vinckeia, Ovale, and the primate-infecting clade containing *P. vivax*-like species), each clustering coherently in the t-SNE space. A single gray un-classified point falls within the Vinckeia cluster (rodent malaria parasites including *P. berghei*, *P. yoelii*, *P. chabaudi* and *P. vinckei*): this corresponds to a genome from the genus *Hepatocystis*, a sister lineage to *Plasmodium* parasitizing bats and Old World monkeys, whose proximity to the rodent malaria subgenus is consistent with the established phylogenetic placement of *Hepatocystis* as nested within mammalian *Plasmodium*. The classification rules used to assign *Plasmodium* species to subgenera are provided in the apicomplexa_classification.json file of the GitHub repository https://github.com/sequana/webapp_kozak for reproducibility.

Apicomplexa typically retain the canonical eukaryotic KCS peak at position –3 but exhibit substantial heterogeneity in upstream signal strength across genera, consistent with the lineage-specific transcriptional and translational regulation reported in this clade. Beyond the genome-level KCS metrics, we applied the Kozak Similarity Score (KSS) to assess per-sequence variation in translation initiation strength across individual Apicomplexa coding sequences. Per-CDS KSS distributions reveal striking differences in initiation signal architecture even between genera with comparable genome-level metrics. For example, *Plasmodium falciparum* and *Toxoplasma gondii* exhibit nearly identical mean KSS values, yet their per-CDS distributions are markedly different (Figure 7): *P. falciparum* exhibits pronounced bimodality, whereas *Toxoplasma gondii* displays a unimodal distribution concentrated around the mean. Whether this bimodality reflects two genuine translational regimes or, in part, the application of a mammalian-derived KSS weight matrix to an extremely AT-rich genome remains to be established (see Limitations). The dedicated Apicomplexa dataset enables detailed exploration of host-adaptation-driven variation in translation initiation and KCS patterns, accessible via the KozakExplorer web application.

**Figure 7:**
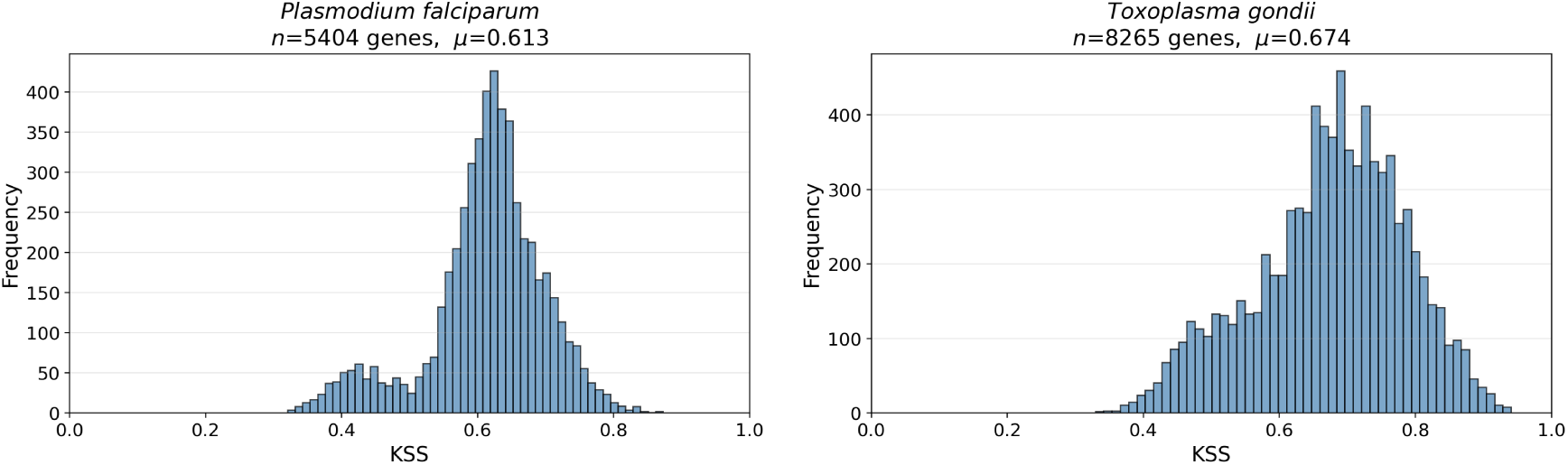
Per-CDS Kozak Similarity Score (KSS) distributions for two Apicomplexa genera. Top: *Plasmodium falciparum* exhibits pronounced bimodality, which may reflect distinct translational regimes or, in part, an artifact of applying the mammalian-derived KSS weight matrix to this AT-rich genome (see Limitations). Bottom: *Toxoplasma gondii* shows a unimodal distribution concentrated around the mean KSS despite having a similar mean KSS value to *P. falciparum*. These contrasting distributions illustrate how per-sequence KSS analysis reveals within-genome heterogeneity that complements genome-level KCS metrics.

### 3.7 Bacterial and archaeal translation initiation patterns

An extended analysis across 22,253 bacterial and 809 archaeal reference genomes provides comparative context for eukaryotic patterns. Bacterial and archaeal genomes exhibit broader, more distributed upstream information patterns consistent with ribosome-binding site architectures, in stark contrast to the highly localized eukaryotic KCS motifs. A comprehensive visualization of global patterns across all domains is provided in the Supplementary Material (Figure S4).

## 4 Discussion

The eukaryotic Kozak landscape and its bacterial and archaeal context together raise several interpretive points that go beyond the descriptive metrics reported above.

### Biological insights

A first salient pattern is the decoupling between positional structure and signal strength. The position of maximum constraint is conserved at −3 across all eukaryotic kingdoms with *n >* 3 except Viridiplantae, where the upstream KCS signal is weak (median KSI ≈ 0.07) and the modal peak instead falls at +4, reflecting the residual G-bias at that position (Table 1). Meanwhile, KSI and *IC_max_*vary by a factor of ∼ 2–3 between lineages (Table 1). Position therefore behaves as a hard structural property of the scanning ribosome and is, in lineages with a resolvable upstream signal, evolutionarily rigid, whereas signal strength is a softer, lineage-tunable parameter reflecting selective pressure on initiation fidelity. This decoupling argues against treating “the” Kozak Consensus Sequence as a single, universally conserved motif and supports a two-axis description that distinguishes *where* the constraint lies from *how strong* it is.

A second salient pattern is the monocot–dicot bipartition within Streptophyta (quantified in the Consistency subsection). That a major angiosperm division emerges unsupervised from initiation-context statistics alone—and reproduces, at far larger scale, differences previously documented only from small species samples [31, 7]—argues that lineage-specific selection has shaped the Kozak context down to sub-kingdom taxonomic resolution, not merely between distant kingdoms.

The dedicated 216-genome Apicomplexa embedding recovers host-range phylogeny: it separates *Plasmodium* sub-genera (Laverania, Vinckeia, Ovale, and the primate-infecting clade) and places *Hepatocystis* inside the rodent-malaria cluster, consistent with its accepted phylogenetic position. Recovering subgenus-level structure from the high-dimensional concatenation of KL, IC, and motif profiles indicates that these per-position vectors carry biological signal rather than only genome-quality covariates. The case study also illustrates how the framework can be redirected to any user-defined clade simply by substituting the input accession list.

Finally, the global cross-kingdom embedding (Figure S4) reveals both domain-level segregation and kingdom-level structure. While eukaryotic, bacterial, and archaeal genomes occupy broadly distinct regions, substantial heterogeneity and overlap (e.g., scattered Chromalveolata genomes) indicate that clustering reflects both phylogeny and lineage-specific biology rather than forming sharply categorical boundaries. While both motifs accurately position the start codon and can be characterized by the same positional-information statistics (KL divergence, information content), they employ distinct recognition mechanisms: direct rRNA–mRNA base-pairing (Shine–Dalgarno) versus nucleotide-context recognition by translation initiation factors (Kozak). This statistical unification does not imply mechanistic equivalence; rather, it demonstrates that diverse biophysical mechanisms produce similar sequence-constraint signatures.

### Methodological considerations

KL divergence is sensitive to the choice of null distribution. KozakExplorer provides three background models—the genomic nucleotide background, the in-frame ATG context, and the shuffled in-frame ATG context—of which the shuffled in-frame ATG background is used for the standardized reference corpus (Supplementary Information). This shuffled control is particularly relevant for GC-rich bacteria, where coding-frame periodicity can otherwise inflate the apparent positional constraint.

The fixed strong/adequate/weak tier scheme [3, 6], although widely used, is a coarse proxy for initiation-context strength and can be misleading when applied across distant lineages. Because it is anchored on only two positions and *requires* a +4 G, it labels every fungal genome as merely adequate even though Fungi carry the strongest and most sharply localized upstream signal in our corpus (Table 2; median KSI 0.187). Conversely, it promotes weakly constrained plant genomes to the strong or optimal tier on the basis of the −3 and +4 positions alone. Continuous information-theoretic summaries (KSI, *IC*) and the per-position profiles avoid this categorical brittleness and are preferable when comparing initiation-context strength across lineages.

t-SNE projections cannot be extended to new genomes without re-fitting. Users analyzing a single new assembly therefore obtain the same per-genome metrics as the reference set, but cannot place their genome directly on the published t-SNE without external re-projection.

All metrics are conditional on annotated start codons, so any misannotation at the level of GFF3 directly biases the upstream context. Conversely, anomalously weak KSI in a clade that otherwise shows a strong consensus may signal potential annotation or assembly quality issues and warrant further investigation, though distinguishing annotation problems from genuine biological divergence requires comparative validation against independent data sources.

The 20/6 window and the canonical-ATG-only restriction are pragmatic defaults. Both are user-tunable in the web application, and a sensitivity analysis is straightforward but currently outside the scope of the standardized landscape.

### Limitations

This study establishes a descriptive geographic boundary for initiation motifs across the tree of life, laying the foundation for future inferential modeling. As a descriptive framework, it is not yet functionally validated against robust experimental datasets (e.g., matched Ribo-seq or polysome profiling) to confirm that higher computationally derived KSI directly predicts higher *in vivo* translational efficiency equivalently across diverse species. Methodologically, KSI is a structurally constrained six-position summary that does not capture longer-range upstream features such as upstream open reading frames or 5’ UTR secondary structures. The per-sequence Kozak Similarity Score (KSS) used in the Apicomplexa case study relies on a weight matrix derived from mammalian cancer transcripts; application to non-mammalian lineages (particularly AT-rich parasitic protists with divergent translation machinery) requires validation against organism-specific or phylogenetically appropriate reference matrices to confirm that observed patterns reflect endogenous biology rather than matrix misalignment. Future statistical modeling of these landscapes will also require formal multiple regression models to rigorously untangle biological variation from potential confounders, such as genome-wide GC content biases and variations in assembly contiguity (N50, total gene count). Finally, the reference analysis is currently restricted to RefSeq-annotated assemblies, leaving understudied lineages under-represented, and adopts a single default background model (the shuffled in-frame ATG pool) for the standardized landscape, so absolute KL and KSI magnitudes should be interpreted relative to that null.

## 5 Conclusion

We introduced KozakExplorer, a comprehensive framework for quantitative analysis of translation initiation signals with primary focus on eukaryotic KCS. Our analysis of 2,282 primary eukaryotic reference genomes reveals a structured yet continuous “KCS landscape” where kingdom-level clustering is evident, peak position remains conserved at –3 across all eukaryotic kingdoms except Viridiplantae (where the weak upstream signal leaves a residual +4 G-bias as the modal peak), and information content varies substantially across lineages. The integration of two comple-mentary information-theoretic profiles (KL divergence relative to background, information content per position) and the derived scalar summaries (KSI, *IC_max_*, *IC_sum_*, peak position) provides a multidimensional characterization of initiation signal architecture.

Two findings carry broader implications. First, classifying genomes by the classical strong/adequate/weak tiers exposes a sharp disagreement with the information-theoretic metrics—most strikingly for Fungi, which combine the strongest, cleanest upstream peak with a mere “adequate” label solely because their consensus lacks a +4 G—demonstrating that fixed, few-position categorical schemes are unreliable proxies for initiation-context strength across distant lineages. Second, the corpus lets us settle a standing question about −3 conservation [7]: a purine occupies the −3 position in 99.2% of eukaryotic genomes, failing only in a handful of highly reduced parasites (chiefly Microsporidia), establishing the −3 purine as a near-invariant anchor of eukaryotic translation initiation.

A dedicated case study of 216 Apicomplexa genomes exemplifies how the framework illuminates host-adaptation-driven structure in parasitic protists, recovering genus-level organization in the Kozak landscape. An extended analysis across 25,344 reference genomes (22,253 bacteria, 809 archaea, 2,282 eukaryota) places eukaryotic patterns within global comparative context, revealing transitions between sharply localized eukaryotic motifs and distributed bacterial and archaeal Shine–Dalgarno signatures.

The `KozakExplorer` web application makes this analysis accessible to researchers across disciplines. By supporting local file uploads, GenBank records, automated NCBI retrieval, and interactive parameter tuning—coupled with real-time visualization and consensus t-SNE embeddings—the tool enables hypothesis generation and annotation quality assessment in both model and non-model eukaryotic organisms. The framework provides a reproducible environment for identifying lineage-specific adaptations and supporting genome annotation refinement.

By integrating standardized information-theoretic metrics with flexible interactive visualization and large-scale em-beddings, `KozakExplorer` establishes a unified and scalable resource for investigating translation initiation signals and KCS across eukaryotic diversity and comparative phylogenetic context.

## 6 Conflicts of interest

The authors declare that they have no competing interests.

## 7 Funding

This work was supported by the ERC Synergy project DecoLeishRN, Grant agreement ID: 101071613.

## 8 Data Availability

The data and source code underlying this article are available in the repositories and under the terms described below.

### Source code and requirements

- **Project name:** KozakExplorer (sequana.kozak module + interactive web application).
- **Project home page:** https://github.com/sequana/webapp_kozak.
- **Source code (library):** https://github.com/sequana/sequana.
- **Operating systems:** platform-independent (tested on Linux and Windows).
- **Programming language:** Python (≥3.10).
- **Other requirements:** pandas, NumPy, SciPy, scikit-learn, Snakemake, Streamlit, logomaker, plotly. A complete list is provided in the pyproject.toml of each repository.
- **Distribution:** PyPI (https://pypi.org/project/sequana/, pip install sequana).
- **License:** BSD 3-clause.

**Pipeline.** The complete Snakemake workflow, together with its analysis parameters (flanking window sizes, filters, background model), is available in the webapp_kozak repository.

**Supporting data.** Genomic sequences and annotations were sourced from NCBI RefSeq and, for the Apicomplexa case study, from a combined RefSeq+GenBank pull. The complete list of accessions and the per-assembly metadata of the Apicomplexa case study are available in the webapp_kozak repository.

**Pre-computed datasets.** All data underlying the t-SNE embeddings and per-genome metrics used in this manuscript are stored as Apache Parquet files in the webapp_kozak repository.

**Repository / DOI.** The code to extract, analyze and quantify Kozak sequences is part of the sequana library (version ≥ 0.22), openly available at https://github.com/sequana/sequana. The Streamlit web application (https://github.com/sequana/webapp_kozak) is currently private and will be made publicly available upon acceptance, together with a Zenodo snapshot of the source code and pre-computed datasets and a citable DOI.

## 9 Author contributions statement

T.C. implemented the Sequana code and the Streamlit application, and wrote the manuscript. A.M.S. proposed the idea and contributed to the writing. J.P.F. contributed to the writing. G.S. contributed to the writing and concept, and funded the study.

## Supplementary Information

This supplementary section provides reference material for KozakExplorer. It covers, in order, the effect of the background model on the KL profile, the auxiliary genome-level scalar metrics omitted from the main text, the global cross-domain embedding, the per-kingdom summary across the full corpus, and the Apicomplexa genus/subgenus classification rules. Each section is self-contained, and the relevant figures and tables are referenced from the main text where they are first discussed.

## S1 Effect of the background model on the KL profile

The KL divergence depends explicitly on the choice of background distribution *P_bg_*(*b*). To illustrate this dependence, Figure S1 shows the per-position KL profile of *Saccharomyces cerevisiae* computed with the three backgrounds exposed in KozakExplorer: the genomic nucleotide background, the in-frame ATG context pool (internal ATG codons, annotated starts removed), and the same in-frame ATG pool with its sequences shuffled per row. All three controls recover the canonical −3 peak—so the location of the constraint is robust—but the peak amplitude varies (from ∼0.35 bits against the genomic background down to ∼0.2 bits against the shuffled in-frame ATG pool). The in-frame ATG pool absorbs codon-position biases of the host genome, lowering the apparent strength of the upstream constraint; the shuffled variant additionally erases positional structure within that pool, providing the most conservative null. For genomes annotated with only a few coding sequences, the in-frame ATG pool may be too small to be reliable, in which case the genome-wide background is preferred.

Both execution paths of KozakExplorer use the same default background: the shuffled in-frame ATG pool (model (iii)). The reproducible Snakemake pipeline that generated the published reference corpus (eukaryota, full cross-domain, and Apicomplexa landscapes; Table 1 and Figure S4) sets `background_method="shuffled"`, and the interactive web application adopts the same default while still exposing the in-frame ATG ("context") and uniform alternatives in its user interface. Because this shuffled null absorbs the codon-frame periodicity that would otherwise contaminate the KL profile of GC-rich genomes, the KL profiles and KSI values reported by the web application are directly comparable with those tabulated for the reference corpus. The location of the inferred constraint is in any case robust to the choice of background (Figure S1); only its magnitude depends on the null.

**Figure S1:**
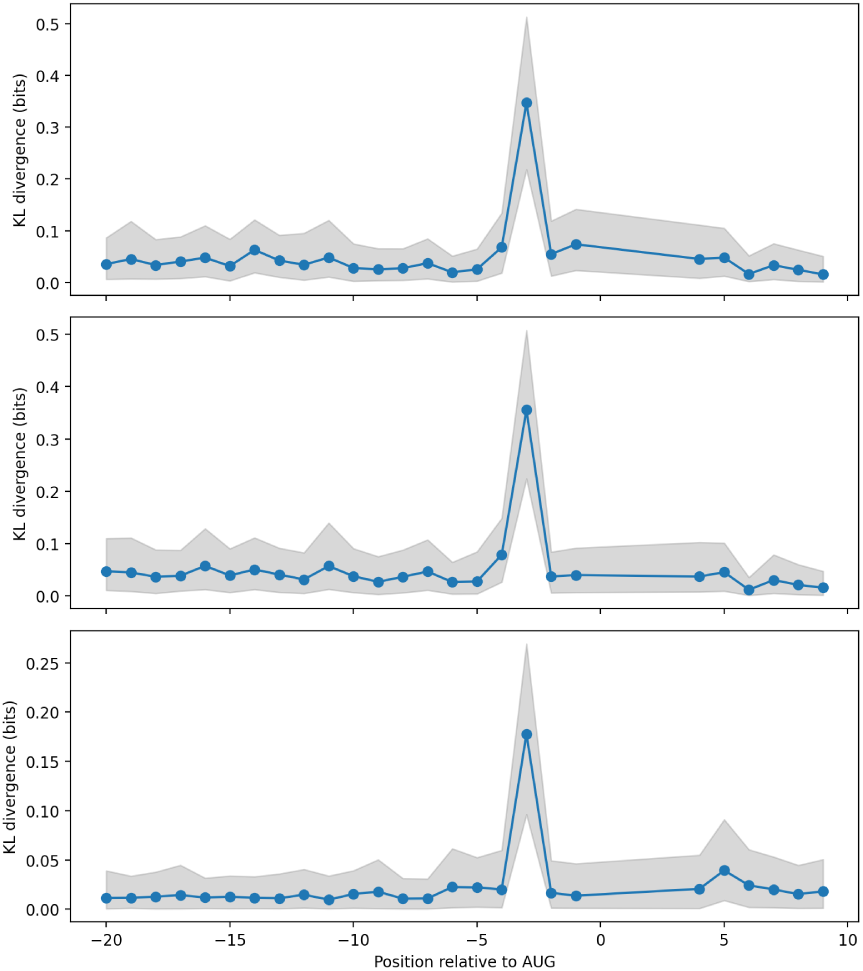
KL divergence profile for *Saccharomyces cerevisiae* (window [−20, +6], ATG starts only) computed with the three background models exposed in KozakExplorer: (top) genome-wide nucleotide background, (middle) inframe ATG pool built from internal ATG codons of the same genome (annotated start sites removed), (bottom) the same in-frame ATG pool with sequences shuffled per row. Shaded bands are bootstrap 95% confidence intervals (200 resamples). The canonical −3 peak is recovered in all three cases; the peak amplitude decreases as the null absorbs more of the codon-position structure of the host genome.

The effect of the background model is most pronounced in high-GC bacterial genomes, where internal ATG codons inherit a strong codon-position bias from the host genome and produce a characteristic 3-bp periodicity. Figure S2 illustrates the three background models on *Streptomyces resistomycificus* (GCF_001514265.1, ∼72% GC). The left column shows the sequence logo of the background pool actually used for each model, and the right column shows the corresponding KL profile of the annotated start-codon contexts against that pool. The genomic and in-frame ATG backgrounds both produce GC-rich pools and yield KL oscillations of ∼0.4 bits at positions −13, −10, −7, reflecting codon-frame nucleotide periodicity rather than a localized biological initiation signal. The shuffled in-frame ATG background, by construction, destroys positional structure within each background sequence: the corresponding logo is visibly more uniform and the KL profile becomes flat across the upstream window, with the residual constraint now concentrated near the start codon (around position −1 to +1). For GC-rich bacterial genomes the shuffled background is therefore the most conservative null and is recommended when the goal is to isolate localized start-codon constraints from codon-frame artifacts; for eukaryotes the difference between the three models is more modest (Figure S1).

**Figure S2:**
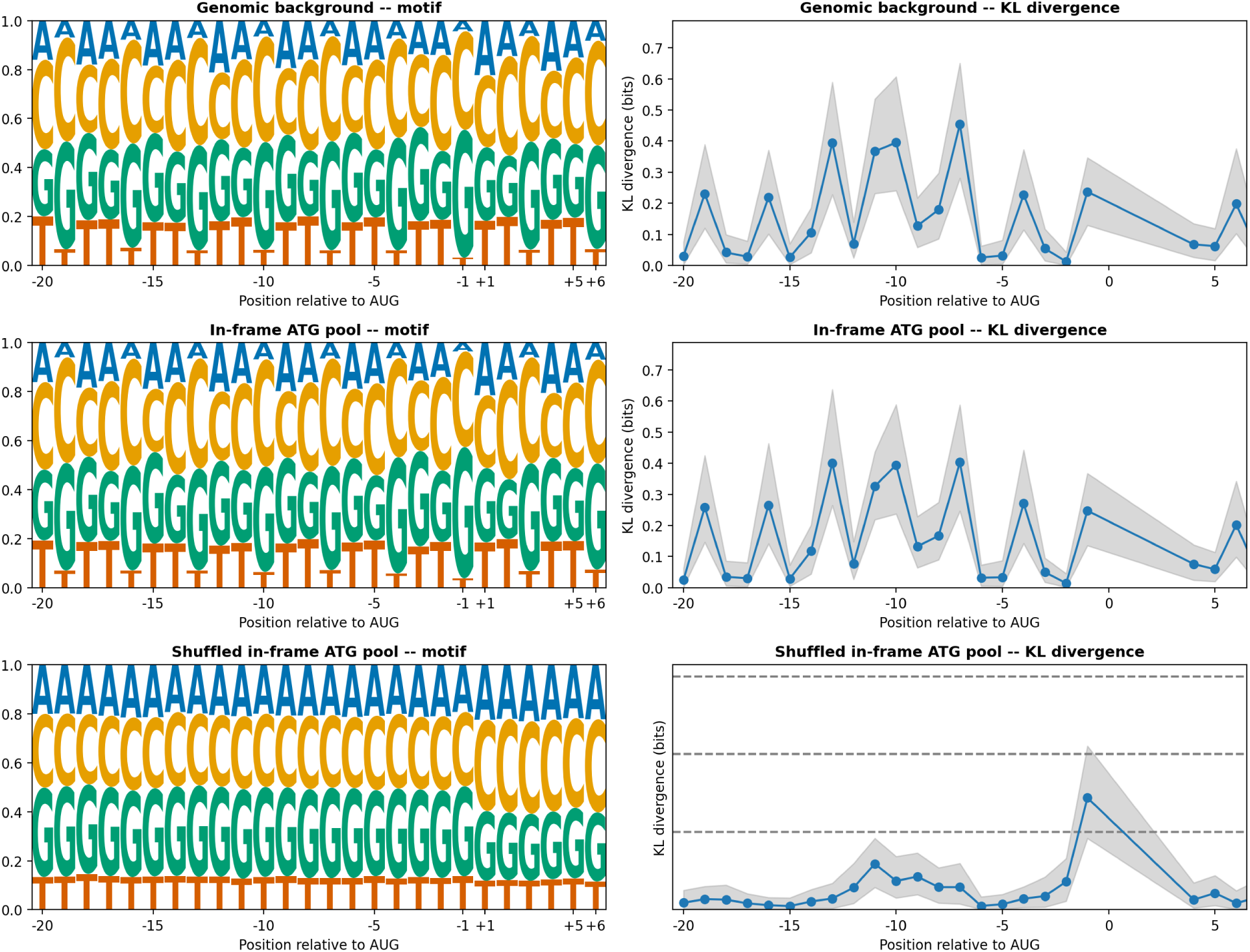
Effect of the three background models on a GC-rich bacterial genome (*Streptomyces resistomycificus*, GCF_001514265.1, ∼72% GC). Rows correspond to the three background models exposed in KozakExplorer: (top) genome-wide nucleotide background, (middle) in-frame ATG pool built from internal ATG codons of the same genome, (bottom) the same in-frame ATG pool with sequences shuffled per row. Left column: sequence logo of the background pool actually used for each model. Right column: KL divergence profile of the annotated start-codon contexts against that pool. The genomic and in-frame ATG nulls retain codon-frame periodicity and yield large ∼0.4- bit oscillations at −13, −10, −7; the shuffled null absorbs that periodicity and leaves a flat upstream KL profile with the residual constraint concentrated near the start codon. Shaded bands: bootstrap 95% confidence intervals (200 resamples).

## S2 Auxiliary genome-level scalar metrics

In addition to the KSI and the information-content metrics (*IC_max_*, *IC_sum_*, position of maximum IC) reported in the main text, the software exports four further scalar summaries derived from the per-position KL profile {*KL_i_*}. These metrics are not used to color the main-text figures and are not part of the t-SNE feature vector, but they are made available for users wishing to characterize a single genome with a richer set of summaries:

- **Total upstream information (***I_total_*): The sum of KL values across the upstream window. 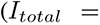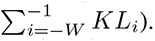. This captures the overall constraint across the upstream window, reflecting the overall intensity of translational constraints.
- **Peak information (***I_peak_*): The maximum positional KL value (*I_peak_* = max*_i_ KL_i_*), which identifies the strongest positional constraint.
- **Signal concentration (***C*): The ratio of peak to total information (*C* = *I_peak_/I_total_*). High *C* values indicate sharply localized motifs, whereas low *C* reflects broadly distributed signals.
- **Effective half-information width (***W*_50_): The minimal number of positions required to account for 50% of *I_total_*. Formally, if the KL values are sorted in descending order:

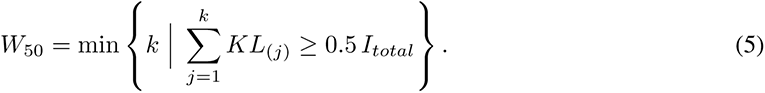

This metric characterizes the effective width of the signal, capturing motif localization independently of arbitrary thresholds.

These auxiliary metrics are exported alongside KSI and the IC summaries in the per-genome parquet datasets distributed with the manuscript.

**Figure S3:**
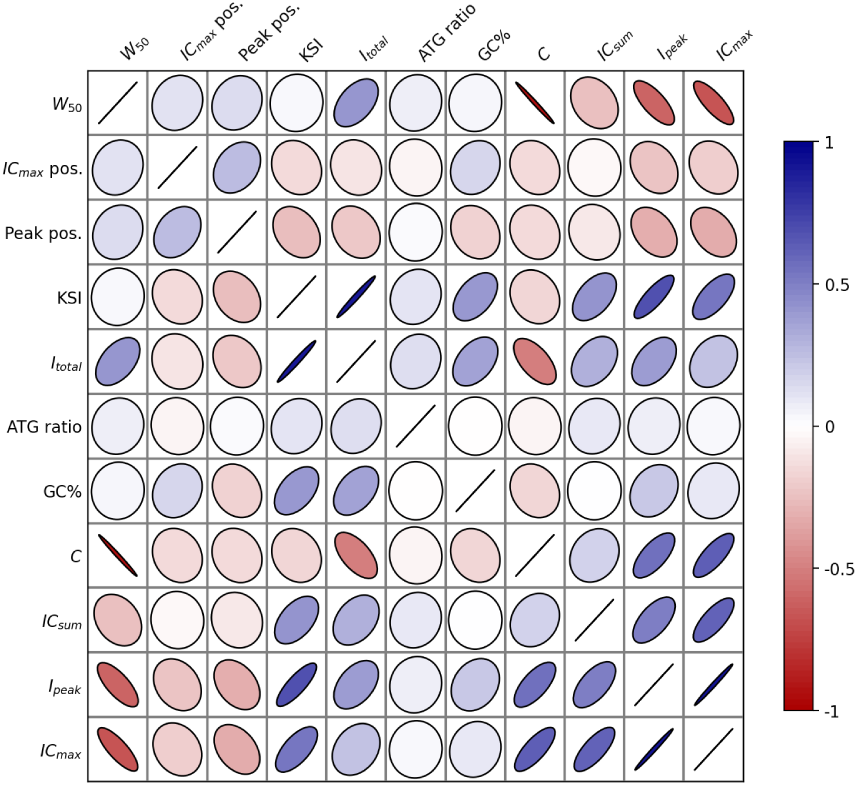
Spearman correlation matrix of genome-level Kozak metrics computed on the 2,282 eukaryotic reference genomes. Ellipse orientation and color encode the sign and magnitude of the rank correlation (blue: positive, red: negative). KSI is strongly correlated with the total upstream information *I_total_*, and *IC_max_* with *I_peak_*, both consistent with their algebraic definitions. In contrast, signal concentration *C*, effective width *W*_50_, peak position, ATG-codon ratio, and GC content show only weak to moderate associations with KSI/IC summaries, confirming that the panel of metrics captures complementary aspects of the initiation signal rather than a single underlying quantity.

## S3 Global landscape of translation initiation signals

The global analysis of 25,344 genomes reveals a highly structured sequence landscape shaped by both phylogenetic descent and GC content pressures (Figure S4). Panel (A) shows that genomes cluster strongly by their domain of life, indicating that the overall signature of translation initiation regions is domain-specific. Panel (B) reveals further organization at the kingdom and phylum levels, with distinct groups corresponding to eukaryotic kingdoms, bacteria, and archaea. The underlying features driving this separation are highlighted in panels (C) and (D). Eukaryotic clusters are dominated by a peak position at −3 (panel C) and moderate total information (panel D), corresponding to localized Kozak motifs. In contrast, bacterial and archaeal clusters exhibit peak positions centered around −11, representing the upstream Shine–Dalgarno interaction site, with total information levels that vary depending on genomic GC content and taxonomy.

**Figure S4:**
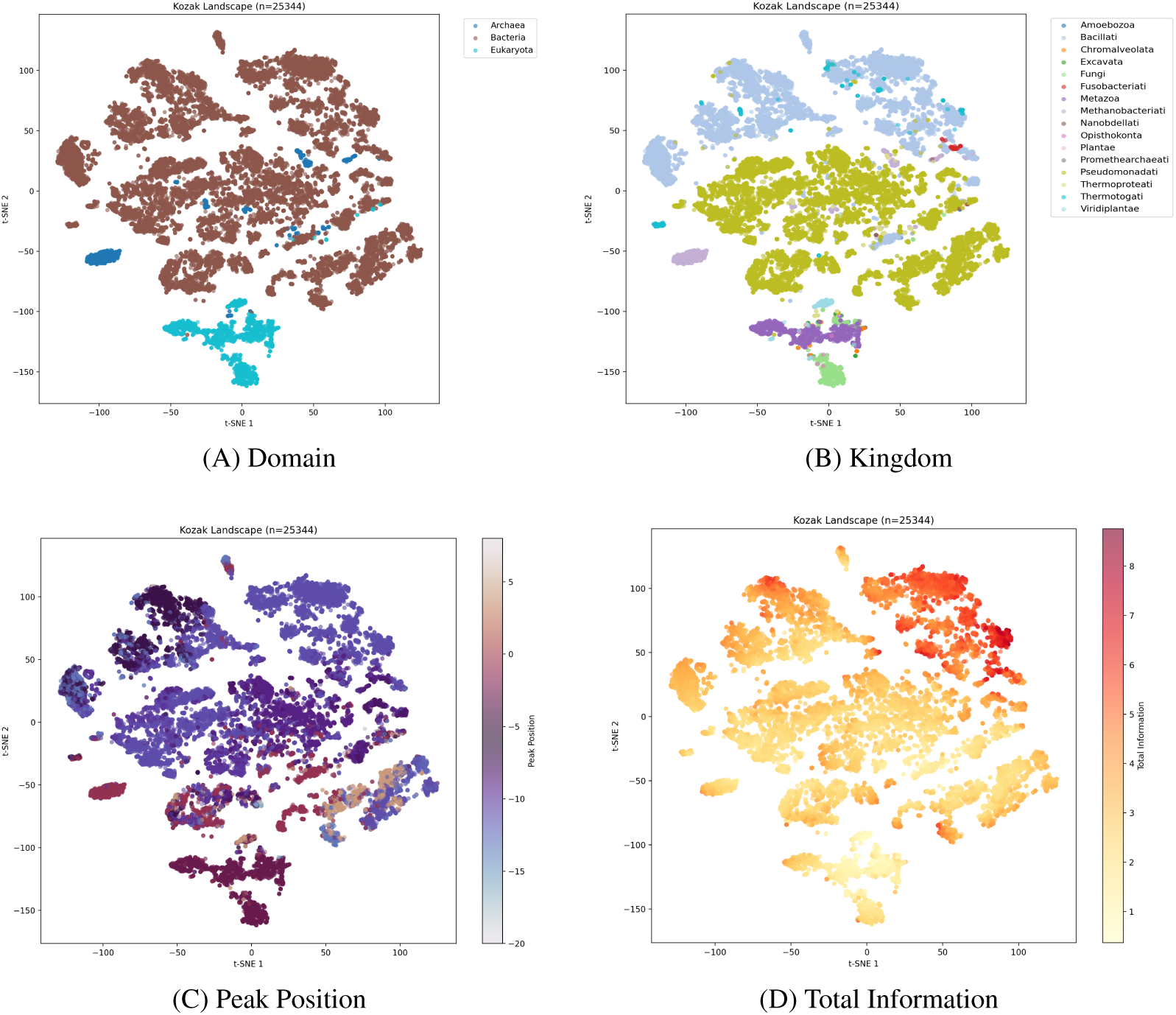
t-SNE projection of genome-level translation initiation signal descriptors for 25,344 reference genomes. (A) Embedding colored by domain of life. (B) Embedding colored by kingdom. (C) Embedding colored by peak position. (D) Embedding colored by total positional information. These visualizations highlight the macroscopic organization of initiation signals across all three domains of life.

**Table S1:**
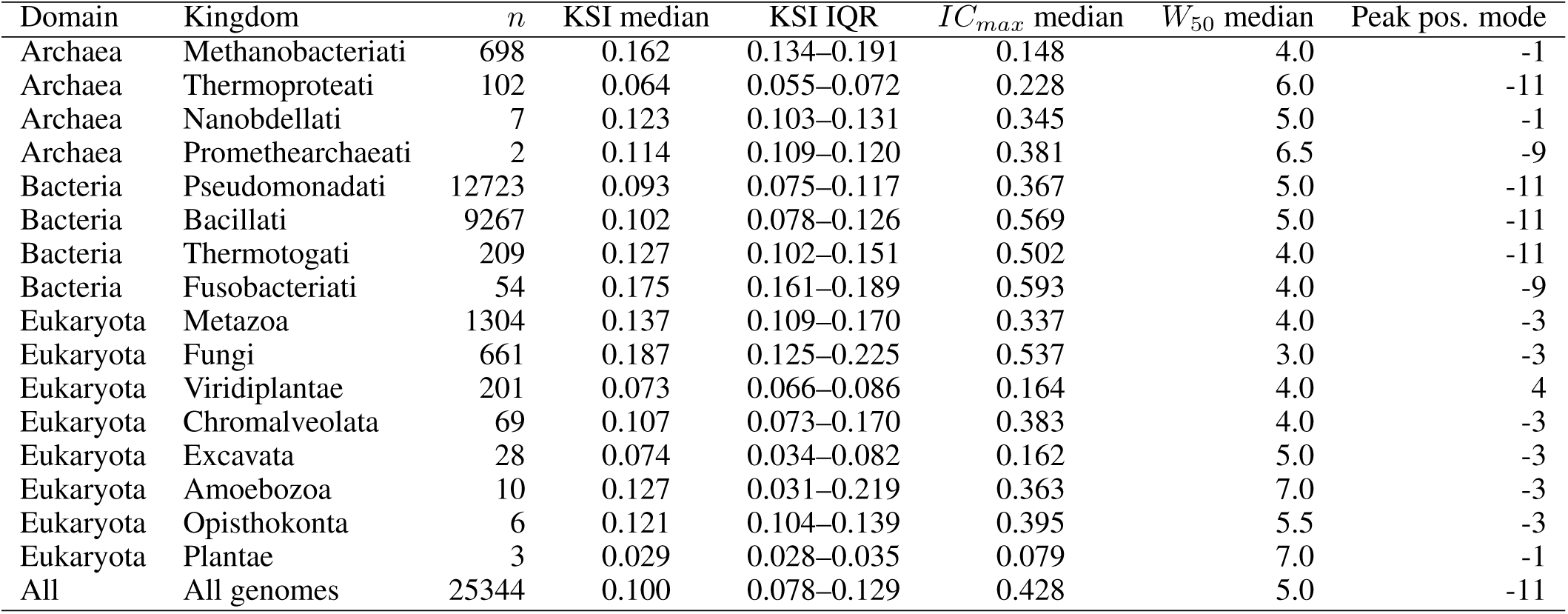
Per-domain and per-kingdom summary of translation initiation metrics across the full reference dataset (Eukaryota, Bacteria, Archaea). Columns are defined as in Table 1.

**Table S2:**
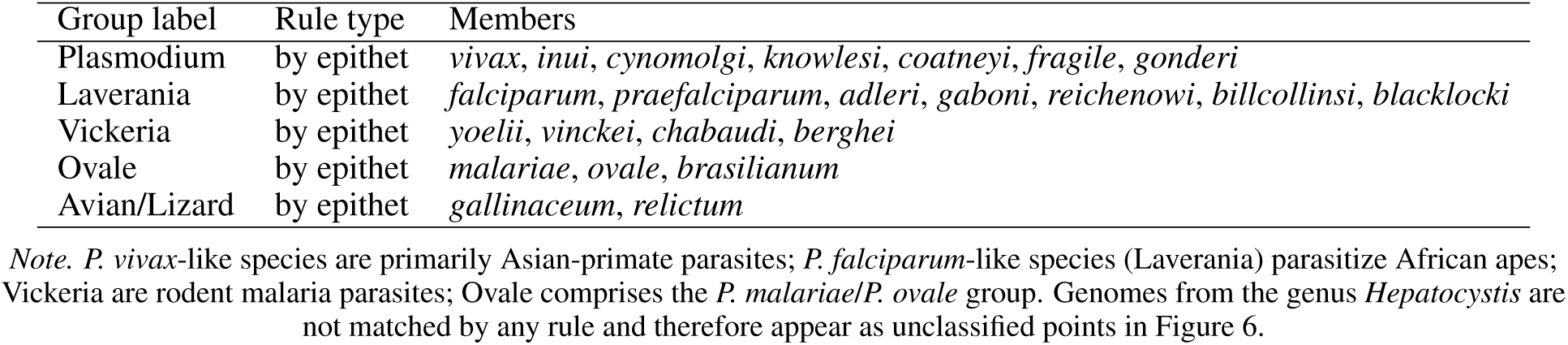
Subgenus and genus assignment rules used to color the Apicomplexa t-SNE embedding (Figure 6). Genus-level rules apply to every species of the listed genus; epithet-level rules map a *Plasmodium* species (by its specific epithet) to a subgenus. Genomes not matched by any rule are shown as unclassified (gray) in the figure. The full machine-readable rule set is provided in `data/apicomplexa_classification.json`.

| Group label | Rule type | Members |
| --- | --- | --- |
| Plasmodium | by epithet | *vivax*, *inui*, *cynomolgi*, *knowlesi*, *coatneyi*, *fragile*, *gonderi* |
| Laverania | by epithet | *falciparum*, *praefalciparum*, *adleri*, *gaboni*, *reichenowi*, *billcollinsi*, *blacklocki* |
| Vickeria | by epithet | *yoelii*, *vinckei*, *chabaudi*, *berghei* |
| Ovale | by epithet | *malariae*, *ovale*, *brasilianum* |
| Avian/Lizard | by epithet | *gallinaceum*, *relictum* |

*Note.* *P. vivax*-like species are primarily Asian-primate parasites; *P. falciparum*-like species (Laverania) parasitize African apes; Vickeria are rodent malaria parasites; Ovale comprises the *P. malariae*/*P. ovale* group. Genomes from the genus *Hepatocystis* are not matched by any rule and therefore appear as unclassified points in Figure 6.

## Notes

### Competing Interest Statement

The authors have declared no competing interest.

